# Reverse engineering morphogenesis through Bayesian optimization of physics-based models

**DOI:** 10.1101/2023.08.21.553928

**Authors:** Nilay Kumar, Alexander Dowling, Jeremiah Zartman

## Abstract

Morphogenetic programs direct the cell signaling and nonlinear mechanical interactions between multiple cell types and tissue layers to define organ shape and size. A key challenge for systems and synthetic biology is determining optimal combinations of intra- and inter-cellular interactions to predict an organ’s shape, size, and function. Physics-based mechanistic models that define the subcellular force distribution facilitate this, but it is extremely challenging to calibrate parameters in these models from data. To solve this inverse problem, we created a Bayesian optimization framework to determine the optimal cellular force distribution such that the predicted organ shapes match the desired organ shapes observed within the experimental imaging data. This integrative framework employs Gaussian Process Regression (GPR), a non-parametric kernel-based probabilistic machine learning modeling paradigm, to learn the mapping functions relating to the morphogenetic programs that generate and maintain the final organ shape. We calibrated and tested the method on cross-sections of *Drosophila* wing imaginal discs, a highly informative model organ system, to study mechanisms that regulate epithelial processes that range from development to cancer. As a specific test case, the parameter estimation framework successfully infers the underlying changes in core parameters needed to match simulation data with time series imaging data of wing discs perturbed with collagenase. Unexpectedly, the framework also identifies multiple distinct parameter sets that generate shapes similar to wild-type organ shapes. This platform enables an efficient, global sensitivity analysis to support the necessity of both actomyosin contractility and basal ECM stiffness to generate and maintain the curved shape of the wing imaginal disc. The optimization framework, combined with fixed tissue imaging, identified that Piezo, a mechanosensitive ion channel, impacts fold formation by regulating the apical-basal balance of actomyosin contractility and elasticity of ECM. This framework is extensible toward reverse-engineering the morphogenesis of any organ system and can be utilized in real-time control of complex multicellular systems.

## 1. Introduction

Reverse engineering biological systems requires mathematical tools to infer interactions and mechanisms of a given process to enable forward engineering applications^1,2^. For instance, a detailed understanding of the pathways involved during a wound healing process can help design new treatment plans or drugs^3,4^. The calibration of computational models of biological systems is challenging due to the large number of interactions whose mathematical description encompasses a very large parameter space. Inevitably, there exists a tradeoff between computational speed and level of detail. For instance, subcellular element models of epithelial morphogenesis can involve hundreds of parameters with an average computational time on the order of days^5^. These models recapitulate a wide variety of biological processes across multiple model organisms with increasing levels of detail. However, before any model can make new predictions, it must be calibrated and validated against experimental data.

On the other hand, in-vivo biological experiments also are costly and time-consuming. The number of features or variables that can be measured in the lab often is limiting. A wide range of forces that often have nonlinear formulations jointly contribute to the shape of an organ. Identifying parameter space that defines similar tissue shapes is the first step in understanding morphogenetic robustness. Moreover, an ability to parametrize a shape also can allow for a more robust comparison between the control, wild-type condition, and a mutant organ shape yielding physical insights (Figure 1). For example, a similar methodology of parametrizing signaling data demonstrates that projecting raw Ca^2+^ signatures from single cells into a more meaningful parameter space of a physics-based model using Approximate Bayesian Computations led to the discovery of four distinct cellular states^6^.

**Figure 1:**
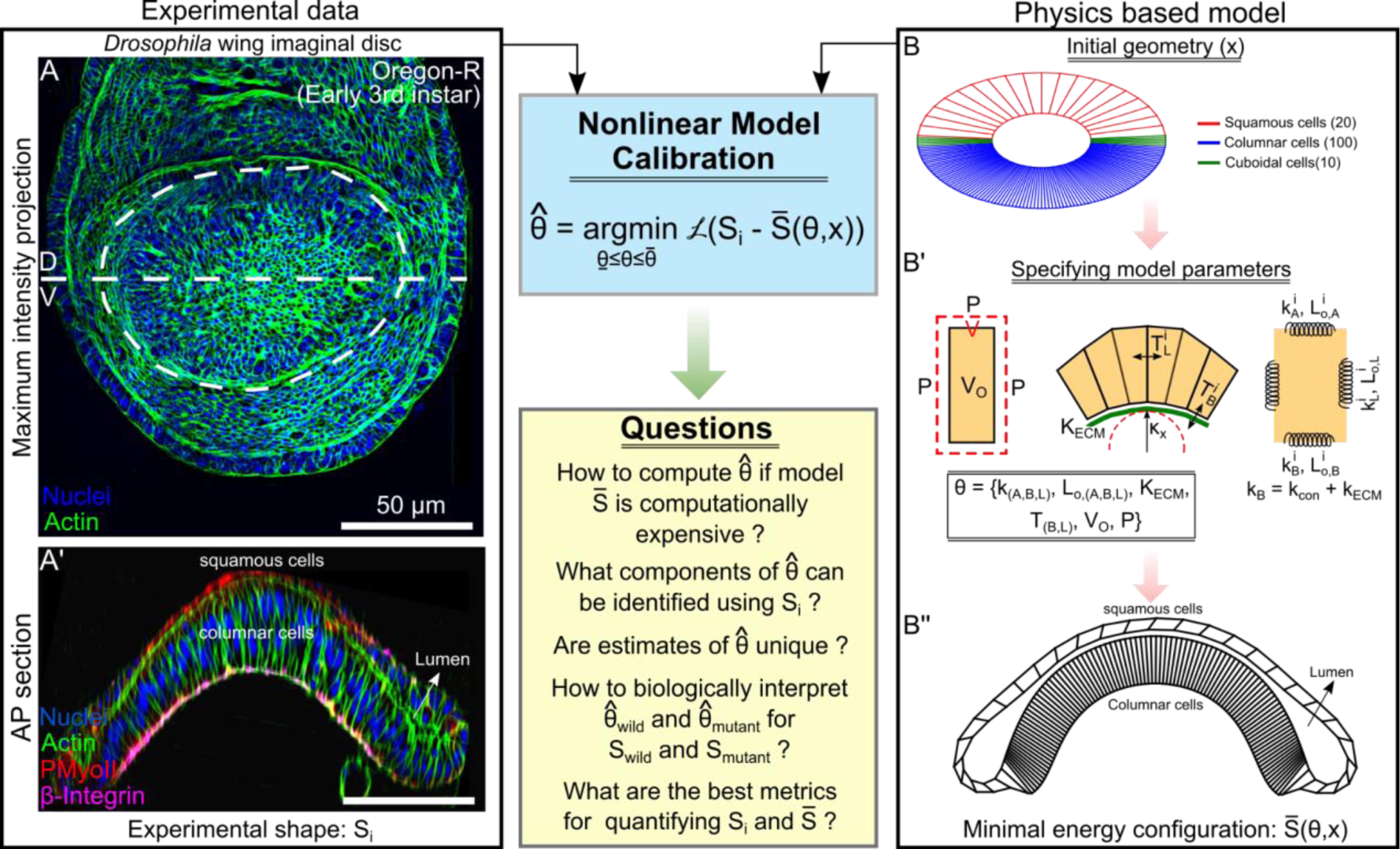
Initial model formulation recapitulates the morphological features of the wing disc. **(A)** Apical view of a z-projection of a *Drosophila* 3^rd^ instar wing imaginal disc. **(A’**) Cross section of the tissue along the anterior-posterior (AP) axis offset from the dorsal-ventral (D, V) boundary. **(B)** The initial geometry used for Surface Evolver simulations. **(B’)** Definition of subcellular cytoskeletal interactions used to define the system’s total energy. **(B’’)** Minimum energy configuration obtained after optimization for parameters in Table 3. Note: The loss function *L* measures the error between the experimental shape *S*_*i*_ and the predicted shape *S̄*(θ, *x*) from the model.

A typical modeling workflow includes the following steps^7^. The overall complex process is conceptually decomposed into a subset of biological processes. These individual subprocesses are first calibrated using a subset of the measurable experimental data before moving towards calibration of the entire process^8^. However, the many interactions between subprocesses often prevent identifying the global optimum. Also, choosing the best error function for comparing experimental data with model output is challenging. Methods of calibrating computational biology models include nonlinear least squares regression^9^, maximum likelihood estimation (MLE), maximum a prior (MAP) estimation^10^, Markov chain Monte Carlo (MCMC)^11^, and genetic algorithms^12^, among others^13–16^.

Each of these algorithms has advantages and disadvantages. While least square and MLE parameter estimation methods typically exhibit fast local convergence, they often get stuck a local minimum using gradient-based optimization methods. MCMC avoids these (sometimes poor) locally optimal parameter point estimates by inferring a posterior distribution, which often requires at least an order of magnitude more computational effort. For instance, MCMC has been used to estimate parameters of ordinary differential equation-based models in systems biology^11^. Another approach followed is Sequential Monte Carlo-Approximate Bayesian Computations (SMC-ABC), where rejection sampling is used to estimate the posterior based on a prior and has been used previously to estimate parameters of models of tissue growth^17^ and Ca^2+^ signaling^6^. However, such an approach is computationally expensive as it requires calculating a distribution and may not be feasible for calibrating computationally expensive modeling frameworks like a molecular dynamic or a subcellular element (SCE) model. Moreover, these detailed modeling approaches require calibration based on multiple measured variables within the lab setup. Consequently, developing new or hybrid approaches that leverage the strengths of multiple approaches can lead to more efficient and robust computational methods. As such, new computationally efficient and robust approaches are needed to calibrate complex mechanistic (biological) mathematical models.

In this work, we present a new computational pipeline employing Bayesian Optimization^18–20^ (BO) to infer the primary biophysical mechanisms driving the shape of an organ. We utilize the *Drosophila* wing imaginal disc cross-section shape (*S*_*i*_) for inferring the parameters (*Θ*) of a biophysical model (*S̄*) of the wing disc cross-section (Figure 1). In general, the framework allows projection of the shape of an organ to a more meaningful parameter space describing biophysical mechanisms driving organ shape generation and maintenance. In this work, we used Surface Evolver^21^ for simulating the wing disc cross-section. Compared to more detailed biological models of wing disc morphogenesis, such as our collaborative SCE model^22^, a model in Surface Evolver simulates fewer details but is less computationally expensive. This modeling framework enables the testing and benchmarking of complex parameter optimization pipelines.

To increase the computational efficiency, the framework couples the mechanistic model to Gaussian process regression (GPR) surrogate models to map the model parameters to the quantitative objective functions while considering uncertainty in model prediction (Figure 2). The surrogate model is then used to sample new points based on an acquisition function that guides the sampling of an optimal solution. This also prevents the model from getting trapped in a local minima. BO is well suited to problems with a large parameter space and easily handles constraints. As such, BO has been applied in a wide range of research areas^23^ including parameter estimation for computationally expensive scientific models^24^. Moreover, several prior studies have demonstrated GPR surrogate models are computationally efficient emulators of biological processes^16^, including post-transcriptional regulation in *Drosophila*^25^, dynamics of microbial systems^26^, cancer tumor growth^27^, and biopharmaceutical manufacturing^28^.

**Figure 2:**
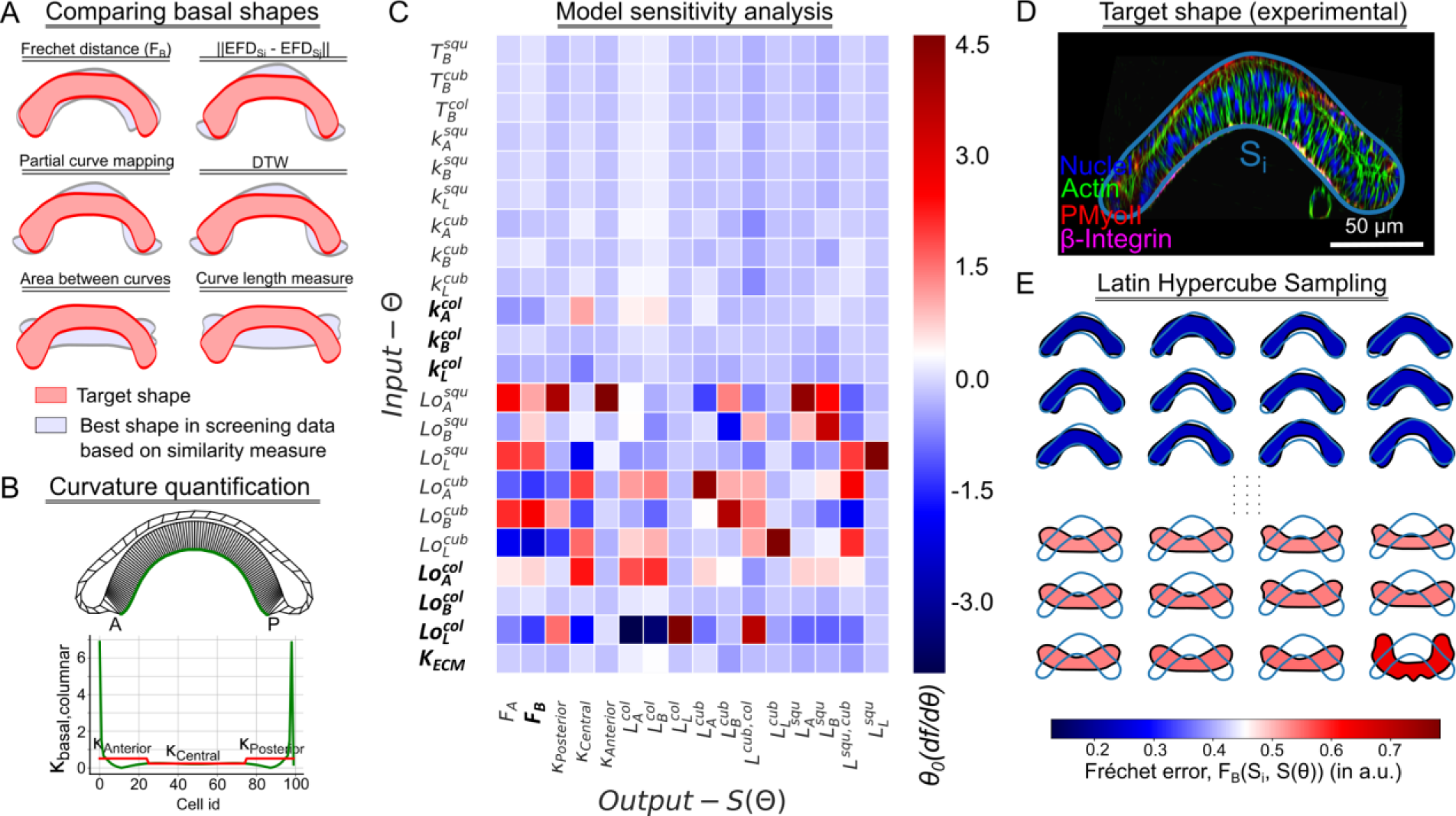
Selection of the optimal quality of fit metric and input parameters for surrogate model training dataset. **(A)** Evaluating similarity measures for computing the objective function. **(B)** Curvature along the basal surface of the columnar cells is plotted as green lines. The basal surface is divided into three equal regions. The average basal curvature is plotted in red for the subregions. **(C)** Heatmap showing a sensitivity measure represented by the color bar legend. The vertical axis represents the input parameter values of the physics-based model, while the horizontal axis represents different morphological features extracted from an output shape. **(D)** A representative AP cross-section of early 3^rd^ instar wing imaginal disc (∼84-90 h AEL), **(E)** Different shapes generated using the parameter sampling are arranged as per increasing Fréchet errors with respect to the representative experimental cross-section.

We first benchmark the pipeline using a synthetic tissue shape with known model parameters (Figure 3). Post benchmarking, we employ the pipeline to predict the morphology of the wing disc undergoing degradation of the extracellular matrix (ECM). To do so, we utilized confocal microscopy data from our earlier investigation^29^ (Figure 4). Lastly, the computational framework, along with fixed tissue imaging data, demonstrated that Piezo, a mechanosensitive ion channel, impacts fold formation within the *Drosophila* wing imaginal disc through the regulation of actomyosin contractility and cell volume (Figure 5). This paves a new direction to discover new mechanisms of mechanotransduction in cytoskeletal regulation during organ growth and morphogenesis. The novel contributions of this work in the domain of systems biology include (1) A successful application of BO to biophysical models of tissue morphogenesis for the first time, (2) a demonstration of Fréchet distance as a useful error metric for calibration of model parameters to define organ shape, and (3) A study highlighting the role of Piezo mediated mechanosensation in fold formation within *Drosophila* wing imaginal disc. Overall, this work provides an efficient pipeline to infer biophysical mechanisms of morphogenesis using morphological data of organs and tissue and identifies a key role of Piezo in regulating tissue shape during wing disc development..

**Figure 3:**
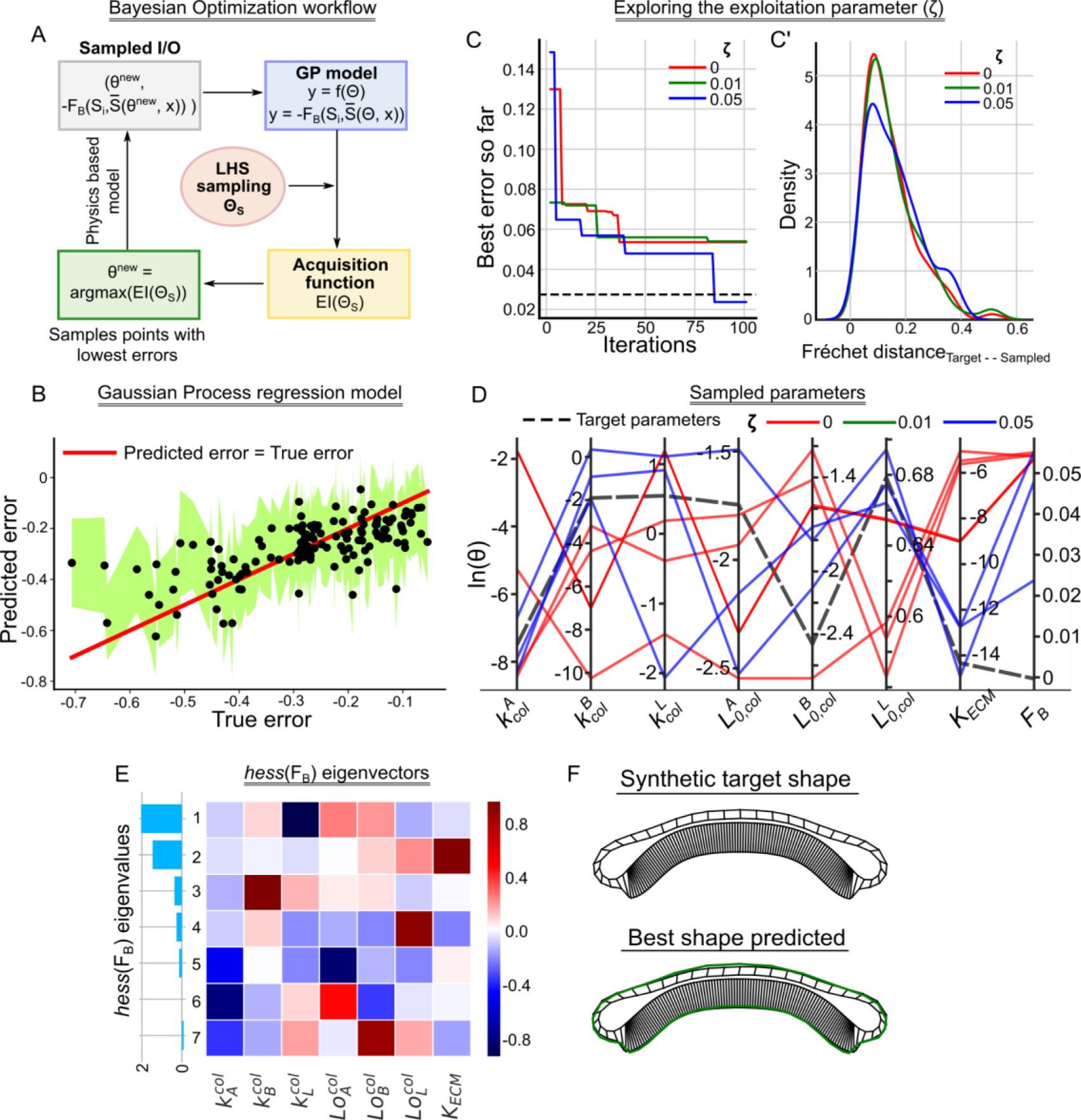
Bayesian optimization using a Gaussian Process regression emulator model enables efficient parameter estimation based on the tissue cross-section shape. **(A)** A schematic for the Bayesian optimization framework. **(B)** A scatter plot of the error predicted by the GPR model against the true error values. The shaded green region represents the lower and upper bound in prediction while the red solid line is a line of parity. **(C)** Plot showing the best error predicted so far with the number of iterations for three different exploration parameter values. **(C’)** Kernel density plots of the error sampled during the BO for three different exploration parameter values. **(D)** A parallel coordinate plot showing the best parameters identified by the BO framework. Different colors represent different *ζ* values and have been included as a legend. Each vertical axis represents one of the parameters represented in a log scale. The last vertical axis represents the F_B_ with respect to the target shape. **(E)** Eigendecomposition analysis of the local Hessian of F_B_. Each row of the heatmap corresponds to the eigenvector of the Hessian matrix. The rows are arranged in decreasing order of the corresponding eigenvalues represented by a bar plot on the left-hand side of the heatmap. **(F)** Tissue geometry for the synthetic target shape and the best shape obtained by the BO framework. The green contour on top of the best shape predicted represents the basal contour of the synthetic target shape.

**Figure 4:**
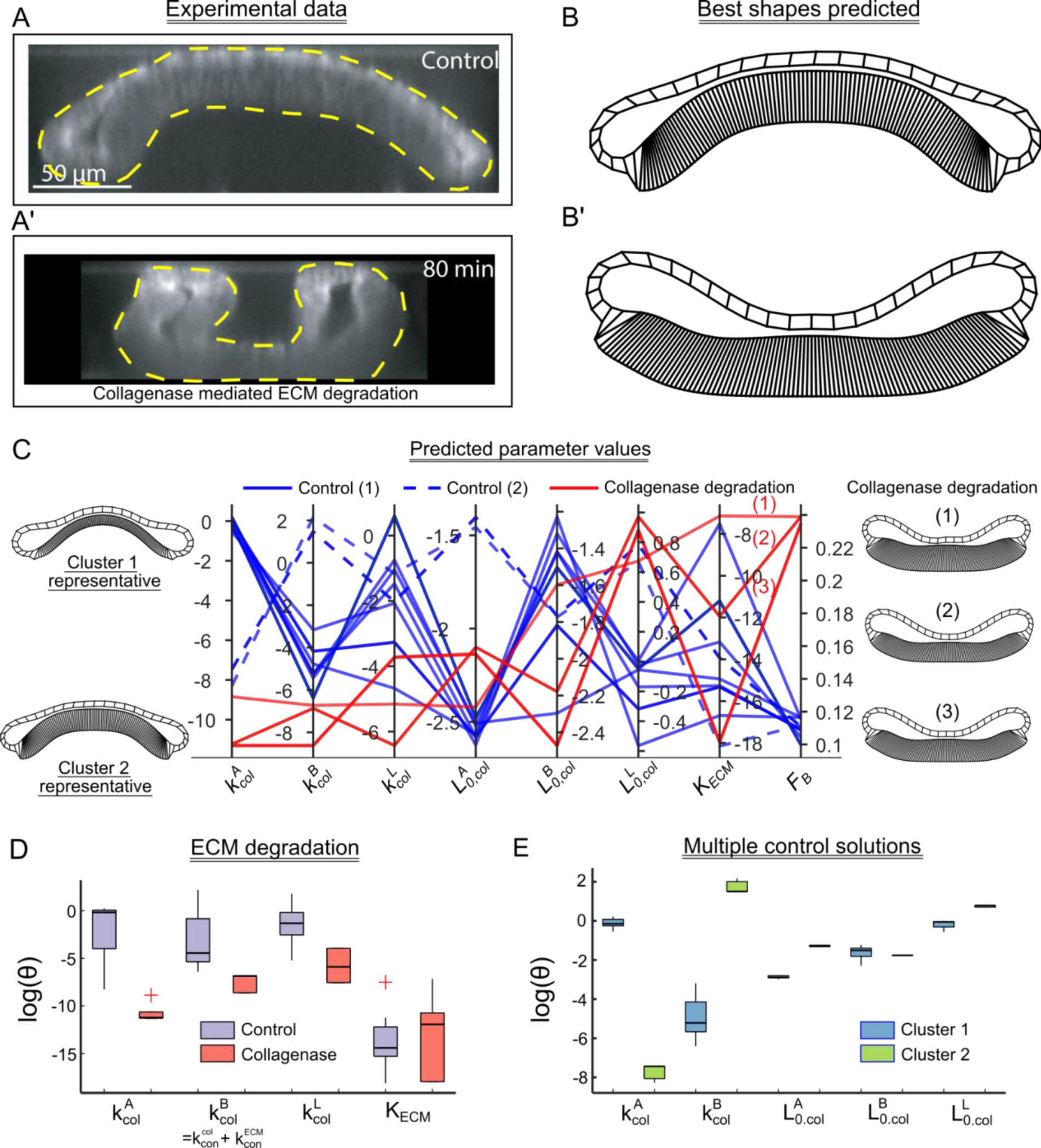
Bayesian Optimization successfully recapitulates the experimental perturbation using only the experimental output shape. **(A-A’)** DV cross-section of wing discs before and after treatment with Collagenase. **(B-B’)** Best shapes predicted by the BO framework in response to the experimental data. **(C)** A parallel coordinate plot showing the best parameters identified by the BO framework. Different colors represent control and mutant samples and have been included as a legend. Each vertical axis represents one of the parameters represented in a log scale. The last vertical axis represents the F_B_ with respect to the target shape. **(D)** Box plot showing the variation of parameters for the best shapes predicted by the BO framework. **(E)** Box plot showing the variation of parameters for the two clusters of parameter sets recapitulating the control “wildtype” shape.

**Figure 5:**
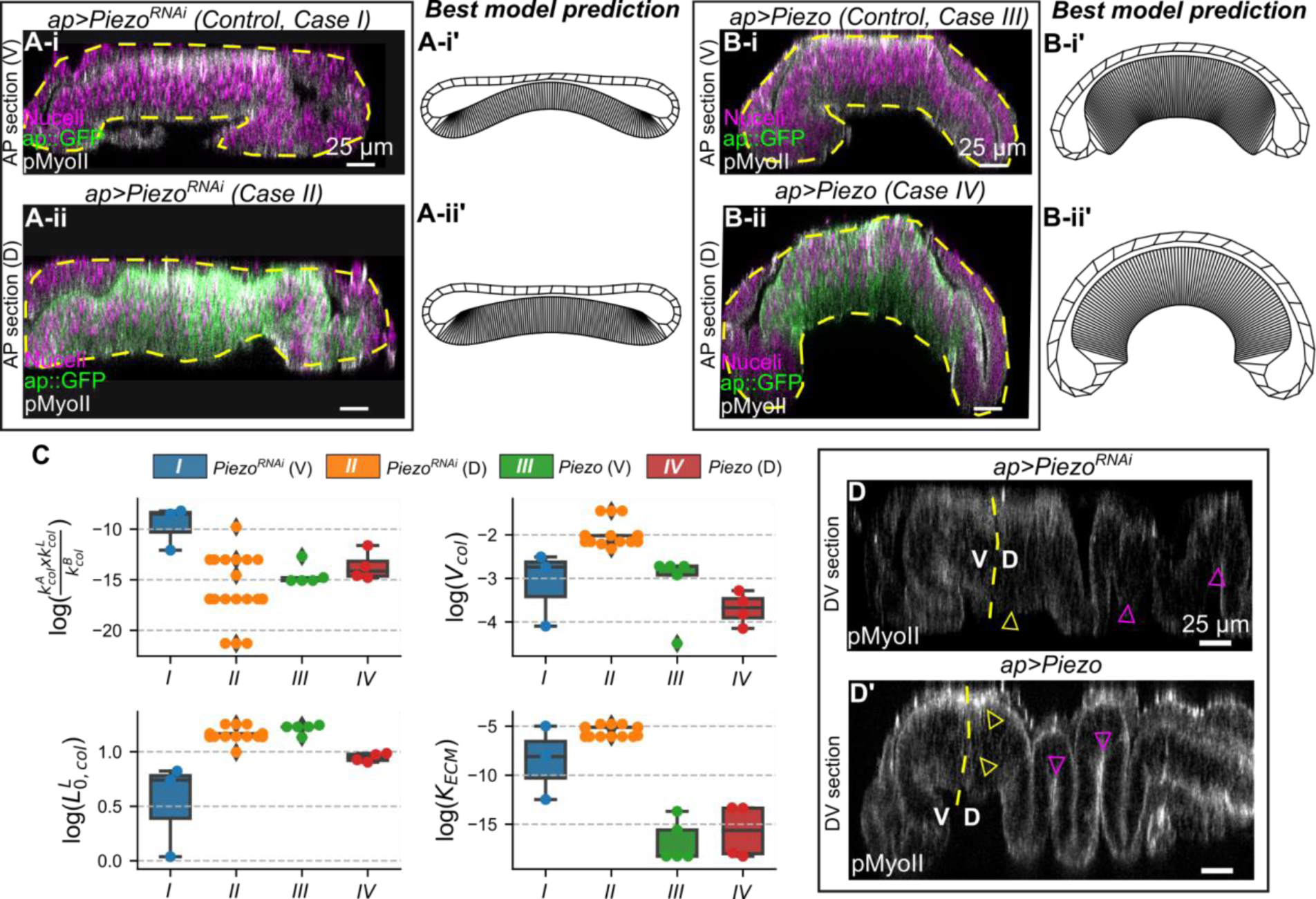
Piezo regulates cell height and fold formation in the *Drosophila* wing imaginal disc. An ap-Gal4 driver was used to downregulate and overexpress Piezo in the dorsal compartment of the wing disc using UAS-Piezo^RNAi^ and UAS-Piezo transgenic fly lines, respectively. **(A-A’)** AP cross sections along the ventral (V, control) and dorsal (D, mutant) compartments of the disc expressing *ap>Piezo^RNAi^*. Fluorescent labels are indicated within the plot. **(A-i’, A-ii’)** Best shapes predicted by the BO framework in response to the experimental data in A-i and A-ii. **(B)** A similar analysis as A for *ap>Piezo*. **(C i-iv)** Box plot showing the variation of parameter combinations (x-axis) for the best shapes predicted by the BO framework. Individual data points have been additionally scattered over the boxes. **(D)** Expression of pMyoII in the DV cross sections of mutant discs. The yellow dashed line represents the approximate dorsal-ventral compartment boundary. Yellow arrows indicate changes in pMyoII and fold formation within the dorsal pouch compartment. At the same time, the magenta arrow suggests changes in the hinge and notum portion of the wing imaginal disc. Please see Figure S4 for additional details..

## 2. Methods and Results

### 2.1 Formulation of a flexible mechanistic model that recapitulates the morphology of the wing disc cross-section

We first utilized Surface Evolver^21^ to formulate a model of the anterior-posterior (AP) cross-section of the *Drosophila* wing imaginal disc (Figure 1A, A’). This approach enabled us to systematically test the utility of benchmarking the Bayesian optimization framework of nonlinear model calibration. The *Drosophila* wing imaginal disc is an established model system for studying the calibration of models of epithelial morphogenesis^30,31^. At later stages of development, the wing disc consists of a fluid-filled sac with a lumen surrounded by epithelial cells of different subtypes (squamous, cuboidal, and columnar) (Figure 1B). A thin extracellular matrix (ECM) encloses the basal surface of cells^32^. The central oval-shaped region termed the pouch resembles a dome-like structure. The shape of the pouch along each direction is patterned by highly conserved morphogens^29,33–35^. The geometrical attributes of wing imaginal disc on a cellular basis was estimated based on a literature review^32,36^ (Table 1), and cell lengths were normalized to avoid numerical instabilities within the model.

**Table 1:**
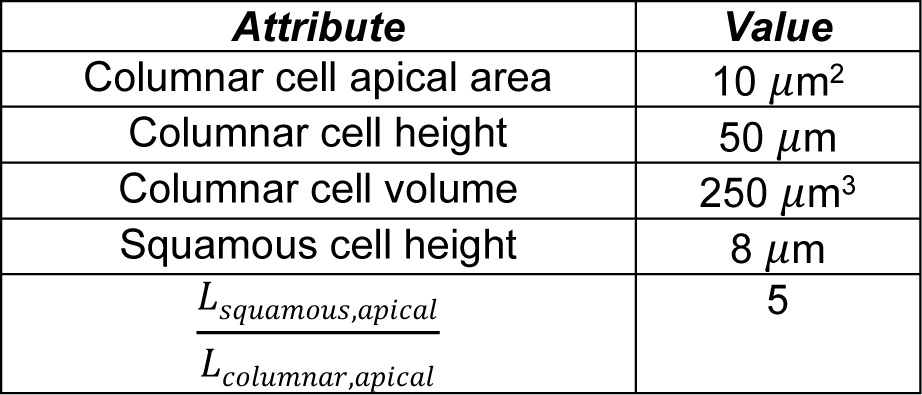
Geometrical attributes of a wing imaginal disc. The quantities reported above are the mean values of the attributes. Note: All cell lengths are normalized against the maximum edge length as surface evolver is known to work best for edges and volumes less than 1.

| **Attribute** | **Value** |
| --- | --- |
| Columnar cell apical area | $10 \mu\text{m}^2$ |
| Columnar cell height | $50 \mu\text{m}$ |
| Columnar cell volume | $250 \mu\text{m}^3$ |
| Squamous cell height | $8 \mu\text{m}$ |
| $\frac{L_{\text{squamous,apical}}}{L_{\text{columnar,apical}}}$ | 5 |

A set of energy functions were defined to incorporate the contributions of known cytoskeletal regulators in the wing imaginal disc (Figure 1B’, Table 2). For instance, phosphorylated non-muscle Myosin II (pMyoII) generates contractile forces by pulling on actin cytoskeleton filaments^37^. This contractility drives shape changes within the tissue that includes fold formation^38,39^. In our model, we assumed each cell edge was a Hookean spring with a natural length, *l_0_*, and a spring constant, *k*. Conceptualizing cell lengths as a spring enables the modeling of length changes. Higher cell contractility increases stiffness and causes resistance to size variations, whereas lower contractility enables more deformations. The energy is calculated using Hooke’s law. Energy is also defined for each individual cell and lumen to penalize changes in volume. The target volume of each component is defined, and Surface Evolver calculates the product of a user-defined constant pressure and any changes in the target volume to estimate the volume energy due to relative compressibility. Apical and lateral adhesion of cells is primarily mediated by apical localization of E-Cadherin and is modeled through the definition of lateral tension^40^. Further, the basal surface of the tissue adheres to the extracellular matrix (ECM) via Integrin adhesion molecules^41,42^. This adhesion is modeled as an additional tension residing in the basal cell edges. Lastly, the ECM is modeled as an elastic string where the energy is evaluated as the integral of the squared curvature over the length of the string. The formulation is similar to one adopted by Storgel et al.^43^. Additionally, the basal contractility (*k*_*B*_) is defined as the sum of actomyosin-mediated contractility at the basal surface (*k*_*con*_) and the ECM (*k*_*ECM*_). We also define a customized repulsion module to stop the apical edges from crossing each other. After every specified number of iterations (50), the repulsion subroutine is called that tracks the distance of each node in the apical surface with the center of all the other apical edges. Suppose the measured distance is less than a particular threshold. In that case, the vertices are shifted away by a minimal distance in the opposite direction normal to the line joining the vertex and the center of the edge.

**Table 2:** Formulation of energy terms included in the Surface Evolver simulations.

|  |  |  |
| --- | --- | --- |
| $E_V$ | Compressibility energy | $P * V$ |
| $E_{\text{spring}}$ | Hookean energy (actomyosin contractility) | $k * (l(e) - l_0(e))^n$ |
| $E_{\text{adhesion}}$ | Cell Tension | $T(e) * l(e)$ |
| $E_{\text{ECM}}$ | Energy of extracellular matrix | $K_{\text{ECM}} \int^{\text{basal}} \kappa^2 dl$ |

Surface Evolver minimizes the system’s total energy under shape constraints to obtain a minimal energy configuration^21^. As a qualitative validation, we chose model parameters based on known experimental constraints (Table 3). Over time, the system’s total energy decreases and then converges to a stable minimum value (Figure S1). This minimum energy configuration qualitatively resembles the experimental cross-section of the wing imaginal disc (Figure 1B’’). The total computational time for convergence of a typical simulation is around 40 minutes using a desktop workstation running Ubuntu 20.04 2 LTS with an Intel® Xeon(R) CPU E5-1603 v3 @ 2.80GHz × 4 processor and 16 GB of RAM. This is ideal for benchmarking a nonlinear model calibration framework as the model provides sufficient detail to capture salient features of the cross-sectional shape but is not too computationally expensive to preclude analysis of the overall pipeline. In the next sections, we describe the sensitivity of model parameters and a framework to compare the error between the model-generated shape with the experimental cross-section.

**Table 3:** Model parameters: The italicized parameters are to be defined within the function that writes the initial geometry file. The other parameters are to be declared within the data file that is used to run Surface Evolver.

| **Parameter name** | **Definition** | **Value** |
| --- | --- | --- |
| $N_{\text{squamous}}$ | Number of squamous cells | 20 |
| $N_{\text{cuboidal}}$ | Number of cuboidal cells | 10 |
| $N_{\text{columnar}}$ | Number of columnar cells | 100 |
| $\text{Pressure}$ | Pressure of system | 0.001 |
| $V_{\text{lumen}}$ | Volume of lumen | 15 |
| $V_{0,\text{cells}}^{\text{sq,cub,col}}$ | Target volume of individual cell type | (0.6,0.6,0.36) |
| $T_{\text{sq}}^{\text{api,bas,lat}}$ | Tension of edges of squamous cells | 0 |
| $T_{\text{cub}}^{\text{api,bas,lat}}$ | Tension of edges of cuboidal cells | 0 |
| $T_{\text{col}}^{\text{api,bas,lat}}$ | Tension of edges of columnar cells | 0 |
| $k_{\text{sq}}^{\text{api,bas,lat}}$ | Spring constant of edges of squamous cells | (0.1,0.1,10) |
| $k_{\text{cub}}^{\text{api,bas,lat}}$ | Spring constant of edges of cuboidal cells | (0.1,0.1,0.1) |
| $k_{\text{col}}^{\text{api,bas,lat}}$ | Spring constant of edges of columnar cells | (0.1,10,0.1) |
| $L_{0,\text{sq}}^{\text{api,bas,lat}}$ | Hooke length of edges of squamous cells | (1,1,0.6) |
| $L_{\text{cub}}^{\text{api,bas,lat}}$ | Hooke length of edges of cuboidal cells | (0.6,0.6,0.6) |
| $L_{\text{col}}^{\text{api,bas,lat}}$ | Hooke length of edges of columnar cells | (0.2,0.2,3) |
| $k_{\text{sq,cub}}$ | Spring constant at cuboidal -squamous interface | 0.01 |
| $k_{\text{col,cub}}$ | Spring constant at cuboidal -columnar interface | 0.01 |
| $L_{\text{sq,cub}}$ | Hooke length of cuboidal-squamous interface | 0.6 |
| $L_{\text{col,cub}}$ | Hooke length of cuboidal-columnar interface | 1.8 |
| $n$ | Power law constant for spring energy model | 2 |
| $K_{\text{ECM,basal}}$ | Modulus of squared curvature energy at ECM and basal cells | 0.01 |
| $K_{\text{apical,lumen}}$ | Modulus | 0.001 |

- sq:squamous, cub:cuboidal, col:columnar, api:apical, bas:basal, lat:lateral

### 2.2 Defining the input-output data for the surrogate model

#### 2.2.1. Fréchet error is the best metric for comparisons between tissue shapes

Here, we first describe a methodology of comparing the outer (basal) contours of any two-wing disc cross sections. The input to a model of the wing disc cross section is the set of parameters (θ) representing cytoskeletal regulation (Figure 1B’). For model calibration, an objective function comparing the target experimental shape (*S*_*i*_) and Surface Evolver generated cross-section (*S̄*(θ, *x*) where: *x* represents the initial geometry) needs to be defined (Figure 1). Elliptic Fourier Descriptors (EFD)^44^ are used to normalize the basal surface of epithelia against the size and translation given the experimental data, and the simulated cross sections are of arbitrary length scales.

To select the best similarity measure for comparing two contours in general, we employed the following strategy: A random simulation was selected from the parameter screening data, and an error was computed based on several similarity measures with respect to the other cross-sections generated within the screen. In particular, we tested the following metrics for comparing two cross-sectional shapes (Figure 2A):

- Area between curves, which is used to compute the area enclosed between any two curves.
- Curve length measure, which is a metric that is used to compare two curves based on their total arc length.
- Elliptic Fourier Descriptors (EFD), which are the calculated Fourier coefficients of the chain-encoded closed contour. This shape descriptor is both rotation and translation invariant. An L2-norm between the EFD coefficients of two shapes is used to quantify similarity^44^.
- Partial Curve Mapping, which is a method of comparing curves through alignments of the smaller curve regions.
- Dynamic Time Warping (DTW), which compares similarities between two signals. It works by distorting one of the signals to maximize its alignment with the other signal with which it is compared^45^.
- Fréchet distance, which is defined as the shortest distance between any two curves given one, is allowed to traverse along the two curves with different speeds^46^.

The best shapes identified using each of the similarity measures listed above for the randomly selected target shape are plotted in Figure 2A. Qualitatively, Fréchet distance^46^ (*F*_*B*_) performed the best in identifying the basal contour most closely approximating the target shape. One advantage of Fréchet distance is that the measure in itself is an error quantification. This simplifies data-driven modeling as it also serves as the objective function. We also report average apical (*L*_*A*_), basal (*L*_*B*_), and lateral (*L*_*L*_) lengths of each cell subtype along with the average basal curvature for the anterior (*κ*_*Anterior*_), medial (*κ_Medial_*), and the posterior (*κ_Posterior_*) halves of the wing disc (Figure 2B).

#### 2.2.2 Parameter sensitivity analysis identifies cell contractility as a key regulator of tissue shape

A sensitivity analysis was carried out to study the effect of varying θ on overall tissue shape (*S̄*(θ, *x*)) (Figure 2C, Figure S2). Each parameter θ was increased and decreased by 70% of its original value. Morphological features within *S̄*(θ, *x*) were measured, and a central finite difference scheme was used to compute the sensitivity of model parameters. As expected, changing the natural lengths of apical, basal, or lateral edges of the squamous, cuboidal, and columnar cells (*L^squ,cub,col^*_*0*,(*A,B,L*)_) caused the most changes across any of the measured features. A loss in the natural length of the cell often represents the cell’s failure to regulate actin polymerization and depolymerization. Dysregulation of actin polymerization causes severe morphological defects^47^.

A change in the spring constants of either of the apical, basal, and lateral edges of a columnar cell (*k*^*col*^_*A*_, *k*^*col*^_*B*_, *k*^*col*^_*L*_) led to changes in the shape of the tissue. It should be noted that the basal contractility in our model is a sum total of contractility generated by the basal actomyosin complex and the ECM. This agrees with our previous work, where we used a more detailed subcellular element model (SCE model) of a wing imaginal disc to show that the tissue shape is mainly generated by actomyosin contractility and maintained by the ECM^29^. We also found that varying the contractility of squamous cells did not impact overall tissue shape (Figure 2C). This observation confirms a recent report where altering contractility and growth through downregulation of PI3K in the squamous epithelia did not significantly impact the shape of wing imaginal disc^32^.

#### 2.2.3 Gaussian Process Regression surrogate model

To further benchmark the calibration of the nonlinear model, we selected a small subset of the parameters identified through sensitivity analysis. The selected parameters are highlighted in bold along the vertical axis of the heatmap within Figure 2C. Latin Hypercube Sampling was used to sample this reduced parameter space uniformly^48^. Surface Evolver was run for all the sampled points, and the geometrical features were extracted. The sampled parameters and the corresponding morphological features constitute the input-output data for a surrogate model to be used for the optimization task. As an example, the sampled cross-sections were arranged based on an increasing Fréchet distance (top to bottom) (Figure 2E) with respect to the target experimental cross-section *S*_*i*_ (Figure 2D). A low Fréchet error corresponds to a better approximation of the experimental data.

Our work used Gaussian Process Regression (GPR) as a surrogate model^49–51^. GPRs, also known as Kriging models, have a rich history in surrogate-assisted optimization^52^, especially for problems with ten to fifty degrees of freedom. We employed a leave-one-out strategy for training where a new model was trained iteratively, leaving exactly one data point out and using everything else for training^51^. The remaining data points were then used for the assessment of the model. With this strategy, we found that the model predictions are in good accordance with the true output.

Based on this result, we utilized all the data points within the parameter screening for the initial training of the GPR model (Figure 3B). In the following sections, we used the described GPR model to develop a framework for Bayesian optimization of the physics-based model of wing disc morphogenesis.

### 2.3 Bayesian optimization of the Gaussian Process Regression (GPR) model of tissue shape enables parameter estimation of the physics-based models

The Bayesian optimization (BO) approach utilizes and updates a prior belief between the inputs and the outputs used for the calculation of the objective function in the form of surrogate models. The surrogate GPR model approximates the functional values for input as a Gaussian distribution allowing quantification of uncertainty in the form of covariances. This is crucial for the computation of acquisition functions, described later in the text, which is used to sample new points within the parameter space, as shown in Figure 3A.

To benchmark the pipeline of parameter estimation, we generated a synthetic target shape with known parameters, which was not included in the training data. For training a GPR model, Fréchet distance (*F*_*B*_) is first computed for all the samples in the screening dataset with respect to the synthetic target shape. The parameters and the corresponding negative value of Fréchet distances (−*F*_*B*_) act as input and output to our GPR model. BO uses an acquisition function guided by the GPR model to draw a new parameter set that maximizes the response value of the GPR. Since we chose −*F*_*B*_ as our response variable, maximizing it corresponds to finding *Θ* leading to the shape closest to the selected target. We use Expected Improvement as our acquisition function. This allows the sampling of new points around the region that maximizes the output of the surrogate model. The exploration parameter (ζ) within the Expected Improvement defines the amount of exploration during the sampling process. Higher exploration parameters tend to sample points from the regions where the GPR model uncertainty is low instead of sampling guided mainly by the mean value of the surrogate model. The pseudo-code for full pipeline is provided as Algorithm 1. This pipeline was implemented in Python (v 3.8.13) using Surface Evolver (v 2.40), Surrogate Modeling Toolbox (v 1.0.0), GPyTorch (v 1.5.0) and scikit-learn (v 0.24.2).

Three different values of the exploration parameter between 0 and 0.05 were selected to run the BO framework for a synthetic tissue cross-section whose parameter values were already known. We plotted the distribution of Fréchet errors for parameters sampled by the BO framework for the different ζ values. A kernel density was calculated to approximate the distributions (Figure 3C’). The plots reveal that increasing ζ tends to flatten out the distributions of errors between the tissue shapes generated by the sampled parameter and the synthetic target shape. This also suggests that the newly sampled points are farther away from the mean of the distribution as compared to points sampled using lower ζ values.

#### Algorithm 1: Bayesian optimization using GPR surrogate models

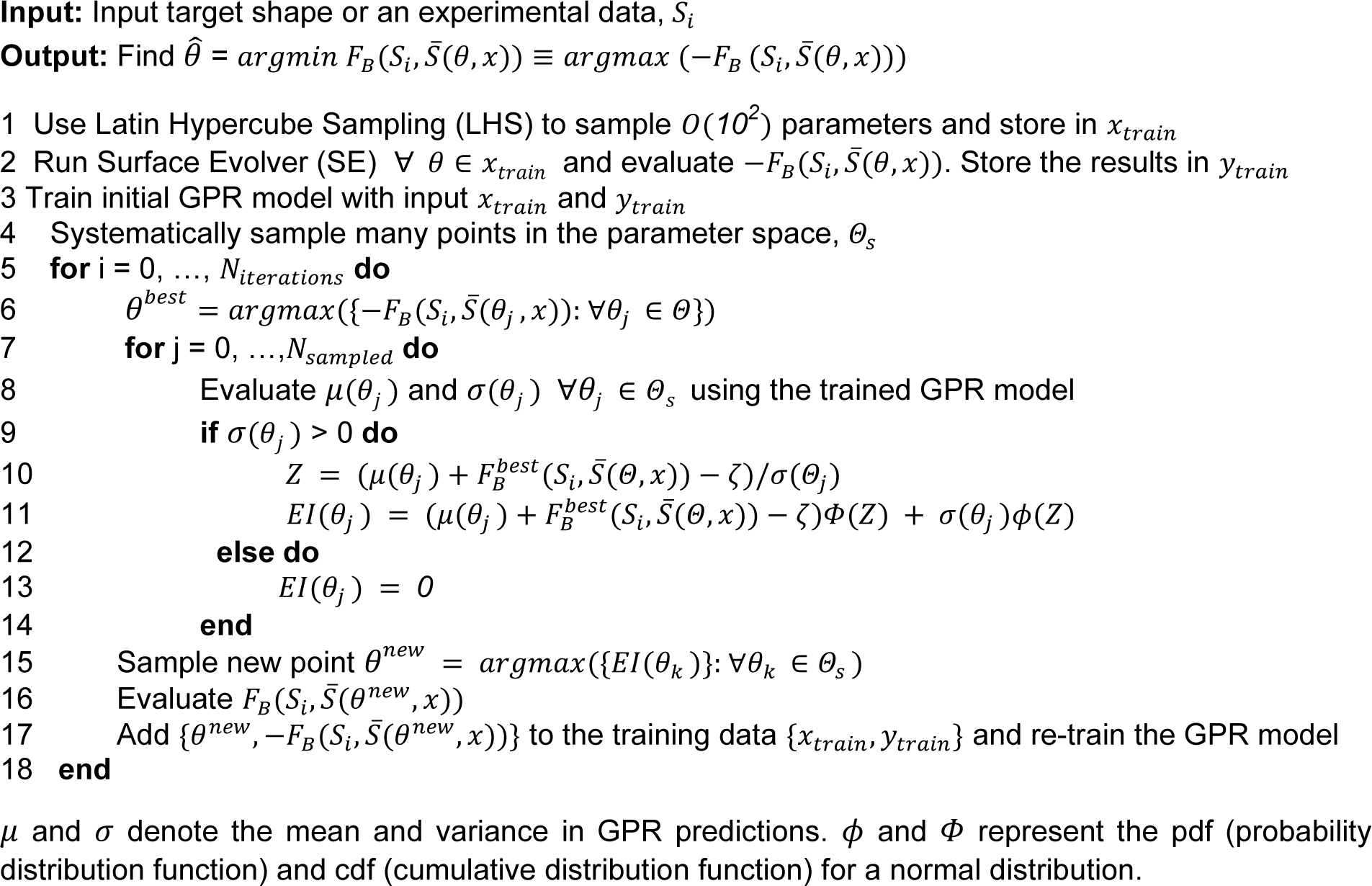

We next plotted the best error sampled so far with each iteration in BO. The best error so far for an i^th^ iteration is defined as the minimal Fréchet distance reported till the particular step during the parameter sampling process. With increasing values of ζ, the points sampled have a lower *F*_*B*_ as compared to *F*_*B*_ from the best shape in training data. An increase in ζ also allows for a faster convergence (Figure 3C). For each ζ, we also plotted the best parameter values that lead to an approximation of the target shape closer or lesser to the best error in the training data. Each vertical axis of the parallel coordinate plot represents the log of parameter values indicated in the horizontal axis label. The dashed black lines represent the true parameter values of the synthetic target shape. The parameters of the best shapes extracted using the pipeline are in good accordance with the synthetic target parameters (Figure 3D, F).

We next analyzed the curvature (Hessian) of the objective function, i.e., F_B_, in the vicinity of the parameters of the target shape to assess the parameters that can be approximated using F_B_ alone (Figure 3E). The eigenvectors corresponding to the two largest eigenvalues predominantly point in the direction of parameters *k*^*col*^_*L*_ and *K*_*ECM*_, respectively. This means that the predicted organ shape, as determined by *F*_*B*_, is most sensitive to these parameters. Thus, *k*^*col*^_L_ and *K*_*ECM*_ can be estimated with the least uncertainty. Previous studies demonstrate that the elasticity of ECM (modeled as *K*_*ECM*_) plays a significant role in determining bulk organ shape^22,32^. A change in lateral contractility (modeled as *k*^*col*^_L_) is also known to cause severe morphogenetic defects as shown by the shape of the cross-section^22^.

The eigenvectors of the third and fourth largest eigenvalues predominantly are in the direction of *k*^*col*^_*B*_ and *L* ^*col*^*_0,L_*, respectively, which means these parameters can be inferred with moderate uncertainty. Finally, the three smallest eigenvalues are near zero, and their eigenvectors predominately point in the direction of *L* ^*col*^*_0,A_*, *k*^*col*^*_A_*, and *L* ^*col*^*_0,B_*. Thus, these parameters are (near) non-estimable using only F_B,_ and the corresponding parameter estimates are highly uncertain.

Previous studies have highlighted the role of basal contractility (modeled as *k*^*col*^_B_) in generating and maintaining the dome shape of wing imaginal disc^29^. A genetic loss of apical-basal contractility through the expression of the dominant negative form of Rho, an upstream regulator of actomyosin contractility, causes the tissue to flatten out^22^. However, a loss of actomyosin contractility upon pharmacological treatment with ROCK inhibitors did not cause severe changes in the shape of a late development stage 3^rd^ instar wing imaginal disc^29^. Genetic perturbation, achieved by expressing Rho^DN^, was performed in the earlier stages of morphogenesis (2^nd^ instar larval stage). In contrast, pharmacological perturbations utilizing ROCK inhibitors were administered during the later larval stage (early 3^rd^ instar larval stage). This suggests that once the wing disc acquires its bent shape, it is less sensitive to changes made in apical-basal actomyosin contractility. Measurements of more variables like local cell lengths can also help better approximate these parameters, which is an open avenue for future investigations.

Through these steps, we conclude that our pipeline employing BO can successfully recover a subset of parameters of the computational model from the representation of the basal surface alone. The proposed methodology allows the transformation of the shape of an organ into a more meaningful parameter space, where each parameter in this space indicates distinct cytoskeletal regulators of wing disc morphogenesis. In the following sections, we describe how to transform mutant organ shapes into the parameter space to identify changes in cytoskeletal regulation that causes changes in the overall tissue shape. Such a pipeline can serve to identify new functions for specific genes and gene products

### 2.4 Bayesian optimization framework predicts loss in contractility of columnar cells upon removal of ECM

As a first test of the framework, Enzymatic degradation of the extracellular matrix (ECM) with Collagenase^29^ was performed to test if the BO framework can recapitulate specific perturbations to cell mechanics, In agreement with our earlier work, the removal of ECM led to a striking inversion or flipping of the curvature within the tissue (Figure 4A, A’), and a loss of inwards bending towards the basal surface of the columnar cell was observed. External contours along the basal surface of the experimental cross-sections are fed into the BO framework as inputs. Our pipeline next estimates the parameters of the Surface Evolver model that would best represent the shape changes. BO was carried out to minimize the Fréchet distance between the experimental cross-section and the simulated shape. The pipeline successfully captured qualitative shape changes as observed within the experimental data as shown in Figure 4B, B’. Significantly, the estimated parameters using the basal contours of the wing disc shape after collagenase treatment indicate a significant decrease in contractility compared to the control (Figure 4C). All apical, basal, and lateral contractility (*k*^*col*^*_A_*, *k*^*col*^*_B_*, *k*^*col*^*_L_*) levels decreased (Figure 4D).

The basal contractility (*k*^*col*^*_B_*) in our model is the sum of contractility imposed by the actomyosin complex (*k*^*col*^*_con_*) and the extracellular matrix contractility (*k*^*ECM*^*_con_*). Modeling these two independently within the Surface Evolver would increase the model complexity. No significant variations were found in the *K*_*ECM*_ parameter, a multiplier of the curvature-based energy used to describe the ECM. The parameter *K*_*ECM*_ contributes towards the elastic energy of the ECM. A higher *K*_*ECM*_ penalizes for higher elastic energy restricting the wing disc from folding, while a lower *K*_*ECM*_ allows wing imaginal disc tissue to make dramatic curvature changes.

We also examined the curvature of the objective function (*F*_*B*_) around the average parameter value obtained from the set of sampled parameters generated by the BO framework (Table S1, Group:Clgns) to assess the identifiability of the model parameters. The Surface Evolver output of the average parameter value yielded a shape similar to one observed when ECM is degraded by collagenase (Figure S3A). Notably, the eigenvector corresponding to the dominant eigenvalue primarily aligns with the parameter *k*^*col*^*_L_* indicating that the predicted organ shape, as described by *F*_*B*_, is most sensitive to variations in the specific parameter (Figure S3B). Consequently, the estimation of *k*^*col*^*_L_* can be carried out with minimal uncertainty. Conversely, the three subsequent eigenvalues exhibit significantly smaller magnitudes than the first one, and their predominant directions align with *K*_*ECM*_, *k*^*col*^*_B_* and *l*^*col*^*_0,L_* respectively. This observation suggests that these parameters can be inferred with moderate uncertainty. Our framework suggests that tissue can still regulate its shape even with varying elasticity by modulating other subcellular features like cell edge lengths and actomyosin contractility. These predictions align with previous findings where a subcellular element model recapitulated the changes upon ECM removal by removing basal contractility and the ECM stiffness from the model^29^.

The control group in our pipeline resulted in two sets (Cluster 1, Cluster 2) of parameters that produced similar tissue shapes (Figure 4C). Examination of the parameters reveals that the differences in shape were due to differences in columnar cell height (*Lo*^*col*^*_L_*) (Figure 4E). Cluster 1 corresponds to tissue shapes with lower lateral cell height but higher apical contractility. In contrast, Cluster 2 corresponds to a higher lateral cell height with higher basal contractility (Table S1). This suggests that maintaining the inwards doming shape with stretching of the pouch cells during growth requires increased basal contractility. It can be achieved either through increasing the ECM stiffness or increasing actomyosin-mediated contractility. Our previous work demonstrated that both ECM stiffness and basal contractility play a crucial role in generating and maintaining the dome shape of the *Drosophila* wing imaginal disc^29^. Other recent work shows increased ECM stiffness as the *Drosophila* wing disc grows in size^32^. Further, the sum of apical and basal contractility is also higher for Cluster 2 than for Cluster 1. Cluster 2 also exhibits cells with lower apical cell area. As the pouch grows in size, the apical cell area decreases in size^36^. The reduction in this apical cell area also contributes to the increase in cell height.

In summary, by applying BO, we show that enzymatic degradation of ECM causes a reduction in global tissue actomyosin contractility. It also causes an inversion in the tissue curvature. Interestingly, our pipeline also reveals two distinct mechanisms of cytoskeletal regulation leading to similar tissue shapes defined by its outer surface. Analysis of the parameters of the two groups further describes biophysical mechanisms driving the thickening of the columnar cells as the tissue grows in size. Our model predicts that an increase in basal contractility and apical cell area is required to sustain the bent shape of the wing imaginal disc as it grows in size.

### 2.5 Piezo regulates basal epithelial curvature through actomyosin contractility, ECM elasticity, and cell volume

Next, we explored the utility of the BO framework for inferring mechanisms that define new morphological shapes downstream of specific genetic perturbations. To do this, we used the Gal4-UAS system to either knockdown or over-express *Piezo* in the dorsal compartment of the wing imaginal disc using an apterous-Gal4 driver. Piezo proteins are a class of mechanosensitive ion channels involved in regulating different biophysical processes^53,54^. However, its role in regulating overall organ shape is poorly understood. Cross-sections parallel to the AP axis were taken in the dorsal side of the pouch guided by the apterous::GFP expressions. The cross-section in the ventral side was used as an internal control. Knockdown of *Piezo* caused loss of doming within the pouch (Figure 5A-ii) along with a reduction in curvature of folds in the notum region of the tissue as compare to the control (Figure S4). On the other hand, overexpression of *Piezo* caused the tissue to increase in bending as compared to the internal control (Figure 5B-ii).

We used the definition of the basal surface of the contours, as shown in Figure 5A, B, as our inputs to the BO framework. Our framework identified shapes qualitatively matching the mutant cross-section (Figure 5A’, B’), A comparative analysis between the model parameters of the best cases identified revealed a decrease in the ratio of apicolateral to basal contractility (defined by *k*^*col*^*_A_* × *k*^*col*^*_L_*/*k*^*col*^*_B_*) upon *Piezo* knockdown as compared to prediction for the internal control. The quantity increased upon overexpression of *Piezo* in a compartment-specific manner (Figure 5C). As predicted through model analysis, quantifying the expression of pMyoII within the *Piezo* mutants also shows an increase in both apical and lateral pMyoII upon overexpression of *Piezo* (Figure 5D’, Figure S4). We also report a decrease in pMyoII in a compartment-specific manner upon *Piezo* knockdown (Figure 5D, Figure S4).

Apart from changes in actomyosin contractility (*k*^*col*^*_i_*), our model also predicts a compartment specific increase in both cell volume (*V*_*cell*_ = *LO*^*col*^*_A_* × *LO*^*col*^*_B_* × *LO*^*col*^*_L_*) and *LO*^*col*^*_B_* upon a knockdown of *Piezo* (Figure 5C). It further predicts a compartment-specific decrease but a global increase in *LO*^*col*^*_L_* on Piezo overexpression as compared to Piezo^RNAi^-expressing discs. An increase in lateral pMyoII levels upon Piezo overexpression can cause the lateral pouch to contract, decreasing its length (Figure 5D’, Figure S4). Interestingly, our pipeline also predicts an overall decrease in the elasticity of ECM upon *Piezo* overexpression (Figure 5C). A previous study of cell dissemination conducted in *Drosophila* midgut has shown that Piezo is required for the degradation of the ECM through Ras^V12^ ^55^. Further experiments are needed to measure changes in ECM elasticity upon Piezo overexpression.

Overall, this section used a combination of quantitative image analysis of fixed tissues and the BO platform to infer potential mechanisms of fold formation regulation by Piezo mechanosensitive ion channels. Our work highlights that *Piezo* regulates bending in *Drosophila* wing imaginal disc by regulating actomyosin contractility, ECM elasticity, or regulation of single cell volume. A more detailed experimental analysis is required to establish mechanisms of cytoskeletal regulation through Piezo.

## Discussion

Morphogenesis involves dynamic shape changes within the organ until it reaches a target size and shape^56,57^. Changes in the global shape of an organ arise from the integration of changes at the cellular level over time. A key challenge for quantitative systems biology is to elucidate how the complex interplay of chemical signals at the single-cell level contributes to the overall organ shape. However, the large diversity of proteins that regulate shape changes and the difficulty of measuring them all at once necessitate the formulation of computational models^58^. A major challenge in calibrating multiscale models of morphogenesis is that they are often highly nonlinear and computationally expensive. Both conventional gradient-based optimization methods (e.g., gradient descent, quasi-Newton methods) and MCMC for Bayesian calibrations require too many evaluations for computationally expensive large-scale models, including subcellular element simulations, to be practical.

In this work, we developed and validated a framework to efficiently infer parameter sets with Bayesian optimization using GPR surrogate models that match experimentally obtained shapes of organs. We found that the optimization process works best using Fréchet distance as the error metric, i.e., the objective function (Figure 2). A multi-faceted Bayesian optimization and local sensitivity analysis revealed that actomyosin contractility and extracellular matrix stiffness are the primary contributors to basal curvature and shape control (Figure 3). This confirms previous reports that relied on experiments and scenarios of more complex subcellular element simulations^29^, demonstrating the robustness of the conclusions.

The current model calculates errors based on overall changes in tissue shape using distances. Often, problems in complex biology are nonlinear and sloppy, leading to multiple parameters within the model that can generate similar shapes^59^. Our pipeline suggests that tissues with different patterns of apical-basal contractility can generate similar shapes of basal curvature while also revealing variable regulation of columnar cell height (Figure 3C). The identification of multiple solutions depends on the number of morphological features used in generating the optimization cost function. For example, including additional components within the objective function in the form of columnar cell height can reduce the number of solutions. The selection rationale of morphological features to include in the cost function depends on the form of experimental data and the choice of the physics-based model.

The BO framework, in conjunction with immunohistochemistry assays, further reveal the role of Piezo mechanosensitive ion channels in regulating fold formation during wing imaginal disc morphogenesis (Figure 5). It does so through a combined regulation of patterning in apical-basal contractility and cell volume. Even though significant efforts have investigated the roles of Piezo in regulating single-cell processes like proliferation, apoptosis, and cytoskeletal regulation^60–64^, emergent functions at the next, multiscale hierarchy of overall organ function remain poorly understood. Previous studies related to Piezo have shown its impact on regulating RhoA^65^. Further, RhoA is upstream of pMyoII, suggesting a potential mechanism. Since a knockdown of Piezo significantly decreased the accumulation of basal pMyoII (Figure 5D, top panel), we also hypothesize that it may be through Integrin clustering mediated activation of mechanosensation. Increased basal contractility reduces the basal cell area bringing integrin molecules sufficiently close to form clusters^66,67^. Once initiated, integrin clusters can stabilize the formation of focal adhesion complexes to increase tension further^68^. The final step can be the activation of Piezo to regulate Rho through Ca^2+^ to further promote pMyoII and consequently contractility^69^. This is equivalent to the autoregulation of pMyoII, which is supported by the experimental evidence that a loss of Integrin in the *Drosophila* wing imaginal disc is also known to reduce basal pMyoII levels^22^. This work thus motivates future experiments to map the exact mechanisms of regulation of pMyoII by Piezo.

Our computational pipeline also predicts a decrease in ECM elasticity (modeled as K_ECM_) upon overexpression of *Piezo* (Figure 5C). Previous studies have proposed the degradation of ECM through Piezo in *Drosophila* midgut^55^. In particular, the loss of Piezo reduced the levels of Mmp2, an enzyme known to degrade ECM. The work further proposed that Piezo can also contribute towards ECM degradation through Ca^2+^-mediated activation of Calpains. Further experimental measurements of ECM elasticity are required to confirm this new predicted function of Piezo during organ development.

Future work can also further define morphogenesis as a multi-objective optimization problem based on multiple outputs of the model and corresponding measurable features in experimental data. The pipeline can be extended for organ-level drug screening, allowing for the study of new mechanical functions of genes due to genetic or pharmacological perturbations. Further, this framework can be extended to studying other physical and biochemical processes such as embryogenesis^70,71^ and models of plant development^72^. Of note, it can also be used to study any models of organogenesis as it is independent of the modeling framework or package used in the physics-based simulation, which makes it attractive for more complex computational models that incorporate subcellular elements^22,29,73,74^. In summary, this computational framework enables the systematic elucidation of generalizable biological rules of the morphogenesis of multicellular systems.

## Methods

### Experimental and image analysis methods

#### Fly stocks and culture

*Drosophila* was grown within an incubator maintained at *25^0^C*. The flies were maintained on 12-hour darkness/light cycle. Virgins from the Gal4 driver colonies were collected twice daily. For the first collection, the bottles are emptied before 6 hours of collection. emale virgins with Gal4 drivers were crossed with UAS-transgene male flies in a 10-15:4 ratio. Early 3^rd^ instar wandering larvae were collected to dissect wing imaginal disc tissue. The wildtype Oregon-R fly line is a long-standing stock in our group originally acquired from the N. Yakoby lab. The following other transgenic stocks and their source, include UAS-Piezo^RNAi^, VDRC # 105132, and UAS-Piezo, BDRC #58772/58773

#### Immunohistochemistry

Wing imaginal discs were dissected in a phosphate-buffered saline (PBS) solution before fixation in a 4% paraformaldehyde in PBS solution. Fixation was done by placing the PCR tubes containing wing disc samples in an ice bath for an hour. Post-fixation, the samples were rinsed with a fresh PBT solution (PBS with 0.03% v/v Triton X-100). Three quick rinses were followed by three 10-minute-long washes. PBT within the tubes was next replaced with 250 μL of 5% normal goat serum (NGS) in PBS and agitated for an hour. Following this, the NGS solution was replaced with the primary antibody solution, and the tubes were left in a rotating platform placed in a temperature of *4^0^C* overnight. The following primary antibodies were used: i) Phospho-Myosin Light Chain 2 (Ser19) (1:50, Rabbit, Cell Signaling Technology #3671S) ii) Integrin *β*PS (myospheroid) (1:5, Mouse, Developmental Studies Hybridoma Bank CF.6G11). The next day, the primary antibody was replaced with PBT. After three quick rinses and three 15-minute long washes, the PBT in the tube is replaced with a secondary antibody solution. The following secondary antibody and dyes were used in our studies: α-Rabbit Alexa Fluor™ 647(1:500, Goat, Thermo Fisher Scientific A32733), α-Mouse Alexa Fluor™ 568 (1:500, Goat, Thermo Fisher Scientific A-11031). DAPI (1:500, Sigma Aldrich D9542). Fluorescein Phalloidin (1:500, Thermo Fisher Scientific F432). After two hours of secondary antibody and dye incubation at room temperature, avoiding light exposure, three rinses of PBT were carried out. After two additional long washes of 15 min, the samples were left for overnight incubation and agitation in PBT at *4^0^C*. The next day, the samples were mounted in a coverslip with spacers to avoid squishing the samples. Spacers were designed using two layers of bio-compatible tapes to create a well to place the samples. Vectashield mounting medium and a cover slip was placed atop, aligned with the spacers.

#### Confocal Microscopy

Imaging of wing imaginal disc samples was done with two different microscopes: Nikon Eclipse Ti confocal microscope with a Yokogawa spinning disc and Nikon A1R-MP laser scanning confocal microscope. For the two confocal microscopes, image data were collected on an IXonEM+colled CCD camera (Andor Technology, South Windsor, CT) using MetaMorph v7.7.9 software (Molecular Devices, Sunnyvale, CA) and NIS-Elements software, respectively. The step size for acquiring 3D data was kept between 0.5-1μm, depending on sample thickness. Imaging was done using 40x and 60x oil objectives with 200 ms exposure time, and 50 nW, 405 nm, 488 nm, 561 nm, and 640 nm laser exposure.

#### Image Analysis

All the raw data presented within the manuscript was analyzed using FIJI/ImageJ. Quantification of fluorescence intensity was carried out using an in-house MATLAB code whose details can be found in the supplementary information of the text. CSBDeep, an ImageJ plugin was used for deconvolution and denoising of the Actin channel. A rolling ball background subtraction was also used to remove background noise. QuickStitch^75^ was used for stitching individual tiles while imaging the entire volume of the imaginal disc.

## Code and data availability

The optimization framework was developed in Python and is available at https://github.com/MulticellularSystemsLab/Tissue-Cartography

## Acknowledgements

NK, and JZ were supported in part by the NIH grant R35GM124935. NK, AD, and JZ were supported by NSF-Simons Center for Quantitative Biology Pilot program and NSF award 2029814. JZ was supported in part by EMBRIO Institute, contract #2120200, a National Science Foundation (NSF) Biology Integration Institute. AD received partial support from NSF Award CBET-1941596. The authors thank P. Brodskiy, A. Madhwan, M.F. Flores, M.S. Mim, M. Unger, V. Velagala, and members of the Multicellular Systems Engineering Lab for their helpful comments and suggestions.

## References

1. Friedel, S., Usadel, B., Von Wirén, N. & Sreenivasulu, N. Reverse Engineering: A Key Component of Systems Biology to Unravel Global Abiotic Stress Cross-Talk. Front. Plant Sci. 3, (2012).

2. Narciso, C. & Zartman, J. Reverse-engineering organogenesis through feedback loops between model systems. Curr. Opin. Biotechnol. 52, 1–8 (2018).

3. Vodovotz, Y. Reverse Engineering the Inflammatory “Clock”: From Computational Modeling to Rational Resetting. Drug Discov. Today Dis. Models 22, 57–63 (2016).

4. Shannon, E. K. et al. Multiple Mechanisms Drive Calcium Signal Dynamics around Laser-Induced Epithelial Wounds. Biophys. J. 113, 1623–1635 (2017).

5. Newman, T. J. Modeling multicellular systems using subcellular elements. Math. Biosci. Eng. MBE 2, 613–624 (2005).

6. Yao, J., Pilko, A. & Wollman, R. Distinct cellular states determine calcium signaling response. Mol. Syst. Biol. 12, 894 (2016).

7. Brodland, G. W. How computational models can help unlock biological systems. Semin. Cell Dev. Biol. 47–48, 62–73 (2015).

8. Anderson, A. E., Ellis, B. J. & Weiss, J. A. Verification, validation and sensitivity studies in computational biomechanics. Comput. Methods Biomech. Biomed. Engin. 10, 171–184 (2007).

9. Reali, F., Priami, C. & Marchetti, L. Optimization Algorithms for Computational Systems Biology. Front. Appl. Math. Stat. 3, (2017).

10. Linden, N. J., Kramer, B. & Rangamani, P. Bayesian parameter estimation for dynamical models in systems biology. PLOS Comput. Biol. 18, e1010651 (2022).

11. Valderrama-Bahamóndez, G. I. & Fröhlich, H. MCMC Techniques for Parameter Estimation of ODE Based Models in Systems Biology. Front. Appl. Math. Stat. 5, (2019).

12. Kumar, M., Husain, D. M., Upreti, N. & Gupta, D. Genetic Algorithm: Review and Application. SSRN Scholarly Paper at 10.2139/ssrn.3529843 (2010).

13. Sun, J., Garibaldi, J. M. & Hodgman, C. Parameter estimation using meta-heuristics in systems biology: a comprehensive review. IEEE/ACM Trans. Comput. Biol. Bioinform. 9, 185–202 (2012).

14. Warne, D. J., Prescott, T. P., Baker, R. E. & Simpson, M. J. Multifidelity multilevel Monte Carlo to accelerate approximate Bayesian parameter inference for partially observed stochastic processes. J. Comput. Phys. 469, 111543 (2022).

15. Warne, D. J., Baker, R. E. & Simpson, M. J. Rapid Bayesian Inference for Expensive Stochastic Models. J. Comput. Graph. Stat. 31, 512–528 (2022).

16. Vernon, I. et al. Bayesian uncertainty analysis for complex systems biology models: emulation, global parameter searches and evaluation of gene functions. BMC Syst. Biol. 12, 1 (2018).

17. Kursawe, J., Baker, R. E. & Fletcher, A. G. Approximate Bayesian computation reveals the importance of repeated measurements for parameterising cell-based models of growing tissues. J. Theor. Biol. 443, 66–81 (2018).

18. Shahriari, B., Swersky, K., Wang, Z., Adams, R. P. & de Freitas, N. Taking the Human Out of the Loop: A Review of Bayesian Optimization. Proc. IEEE 104, 148–175 (2016).

19. Greenhill, S., Rana, S., Gupta, S., Vellanki, P. & Venkatesh, S. Bayesian Optimization for Adaptive Experimental Design: A Review. IEEE Access 8, 13937–13948 (2020).

20. Ulmasov, D., Baroukh, C., Chachuat, B., Deisenroth, M. P. & Misener, R. Bayesian Optimization with Dimension Scheduling: Application to Biological Systems. in Computer Aided Chemical Engineering (eds. Kravanja, Z. & Bogataj, M.) vol. 38 1051–1056 (Elsevier, 2016).

21. Brakke, K. A. The Surface Evolver. Exp. Math. 1, 141–165 (1992).

22. Kumar, N. et al. Balancing competing effects of tissue growth and cytoskeletal regulation during Drosophila wing disc development. 2022.09.28.509971 Preprint at 10.1101/2022.09.28.509971 (2022).

23. Wang, K. & Dowling, A. W. Bayesian optimization for chemical products and functional materials. Curr. Opin. Chem. Eng. 36, 100728 (2022).

24. Befort, B. J., DeFever, R. S., Tow, G. M., Dowling, A. W. & Maginn, E. J. Machine Learning Directed Optimization of Classical Molecular Modeling Force Fields. J. Chem. Inf. Model. 61, 4400–4414 (2021).

25. López-Lopera, A. F., Durrande, N. & Álvarez, M. A. Physically-Inspired Gaussian Process Models for Post-Transcriptional Regulation in Drosophila. IEEE/ACM Trans. Comput. Biol. Bioinform. 18, 656–666 (2021).

26. Oyebamiji, O. K. et al. Gaussian process emulation of an individual-based model simulation of microbial communities. J. Comput. Sci. 22, 69–84 (2017).

27. Rocha, H. L., de O. Silva, J. V., Silva, R. S., Lima, E. A. B. F. & Almeida, R. C. Bayesian inference using Gaussian process surrogates in cancer modeling. Comput. Methods Appl. Mech. Eng. 399, 115412 (2022).

28. Tulsyan, A. et al. Spectroscopic models for real-time monitoring of cell culture processes using spatiotemporal just-in-time Gaussian processes. AIChE J. 67, e17210 (2021).

29. Nematbakhsh, A. et al. Epithelial organ shape is generated by patterned actomyosin contractility and maintained by the extracellular matrix. PLOS Comput. Biol. 16, e1008105 (2020).

30. Beira, J. V. & Paro, R. The legacy of Drosophila imaginal discs. Chromosoma 125, 573–592 (2016).

31. Smith-Bolton, R. Drosophila Imaginal Discs as a Model of Epithelial Wound Repair and Regeneration. Adv. Wound Care 5, 251–261 (2016).

32. Harmansa, S., Erlich, A., Eloy, C., Zurlo, G. & Lecuit, T. Growth anisotropy of the extracellular matrix drives mechanics in a developing organ. 2022.07.19.500615 Preprint at 10.1101/2022.07.19.500615 (2022).

33. Parker, J. & Struhl, G. Control of Drosophila wing size by morphogen range and hormonal gating. Proc. Natl. Acad. Sci. 117, 31935–31944 (2020).

34. Bejsovec, A. Wingless Signaling: A Genetic Journey from Morphogenesis to Metastasis. Genetics 208, 1311–1336 (2018).

35. Teleman, A. A. & Cohen, S. M. Dpp Gradient Formation in the Drosophila Wing Imaginal Disc. Cell 103, 971–980 (2000).

36. Breen, D. E., Sui, L., Bai, L., Jülicher, F. & Dahmann, C. Cell-level 3D reconstruction and quantification of the Drosophila wing imaginal disc. Int. J. Bioinforma. Res. Appl. 15, 174–189 (2019).

37. Betapudi, V. Life without double-headed non-muscle myosin II motor proteins. Front. Chem. 2, (2014).

38. Guo, H., Swan, M. & He, B. Optogenetic inhibition of actomyosin reveals mechanical bistability of the mesoderm epithelium during Drosophila mesoderm invagination. eLife 11, e69082 (2022).

39. Heer, N. C. et al. Actomyosin-based tissue folding requires a multicellular myosin gradient. Dev. Camb. Engl. 144, 1876–1886 (2017).

40. Wodarz, A., Stewart, D. B., Nelson, W. J. & Nusse, R. Wingless signaling modulates cadherin-mediated cell adhesion in Drosophila imaginal disc cells. J. Cell Sci. 119, 2425–2434 (2006).

41. Domínguez-Giménez, P., Brown, N. H. & Martín-Bermudo, M. D. Integrin-ECM interactions regulate the changes in cell shape driving the morphogenesis of the Drosophila wing epithelium. J. Cell Sci. 120, 1061–1071 (2007).

42. Fristrom, D., Wilcox, M. & Fristrom, J. The distribution of PS integrins, laminin A and F-actin during key stages in Drosophila wing development. Dev. Camb. Engl. 117, 509–523 (1993).

43. Štorgel, N., Krajnc, M., Mrak, P., Štrus, J. & Ziherl, P. Quantitative Morphology of Epithelial Folds. Biophys. J. 110, 269–277 (2016).

44. Kuhl, F. P. & Giardina, C. R. Elliptic Fourier features of a closed contour. Comput. Graph. Image Process. 18, 236–258 (1982).

45. Dynamic Time Warping. in Information Retrieval for Music and Motion (ed. Müller, M.) 69–84 (Springer, 2007). doi:10.1007/978-3-540-74048-3_4.

46. Alt, H. & Godau, M. Computing the fréchet distance between two polygonal curves. Int. J. Comput. Geom. Appl. 05, 75–91 (1995).

47. Eaton, S., Auvinen, P., Luo, L., Jan, Y. N. & Simons, K. CDC42 and Rac1 control different actin-dependent processes in the Drosophila wing disc epithelium. J. Cell Biol. 131, 151–164 (1995).

48. Tang, B. Orthogonal Array-Based Latin Hypercubes. J. Am. Stat. Assoc. 88, 1392–1397 (1993).

49. Tonner, P. D., Darnell, C. L., Engelhardt, B. E. & Schmid, A. K. Detecting differential growth of microbial populations with Gaussian process regression. Genome Res. 27, 320–333 (2017).

50. Schulz, E., Speekenbrink, M. & Krause, A. A tutorial on Gaussian process regression: Modelling, exploring, and exploiting functions. J. Math. Psychol. 85, 1–16 (2018).

51. Rasmussen, C. E. & Williams, C. K. I. Gaussian Processes for Machine Learning. (2005). doi:10.7551/mitpress/3206.001.0001.

52. Fuhg, J., Fau, A. & Nackenhorst, U. State-of-the-Art and Comparative Review of Adaptive Sampling Methods for Kriging. Arch. Comput. Methods Eng. 28, (2020).

53. Wu, J., Lewis, A. H. & Grandl, J. Touch, Tension, and Transduction – The Function and Regulation of Piezo Ion Channels. Trends Biochem. Sci. 42, 57–71 (2017).

54. Honoré, E., Martins, J. R., Penton, D., Patel, A. & Demolombe, S. The Piezo Mechanosensitive Ion Channels: May the Force Be with You! in Reviews of Physiology, Biochemistry and Pharmacology Vol. 169 (eds. Nilius, B. et al.) 25–41 (Springer International Publishing, 2015). doi:10.1007/112_2015_26.

55. Lee, J., Cabrera, A. J. H., Nguyen, C. M. T. & Kwon, Y. V. Dissemination of RasV12-transformed cells requires the mechanosensitive channel Piezo. Nat. Commun. 11, 3568 (2020).

56. Heisenberg, C.-P. & Bellaïche, Y. Forces in Tissue Morphogenesis and Patterning. Cell 153, 948–962 (2013).

57. Buchmann, A., Alber, M. & Zartman, J. J. Sizing it up: the mechanical feedback hypothesis of organ growth regulation. Semin. Cell Dev. Biol. 35, 73–81 (2014).

58. Wu, Q., Kumar, N., Velagala, V. & Zartman, J. J. Tools to reverse-engineer multicellular systems: case studies using the fruit fly. J. Biol. Eng. 13, 33 (2019).

59. Goodwin, B. C., Kauffman, S. & Murray, J. D. Is Morphogenesis an Intrinsically Robust Process? J. Theor. Biol. 163, 135–144 (1993).

60. Nourse, J. L. & Pathak, M. M. How cells channel their stress: Interplay between Piezo1 and the cytoskeleton. Dev. Biol. (2017).

61. Tsai, F.-C., Kuo, G.-H., Chang, S.-W. & Tsai, P.-J. Ca2+ signaling in cytoskeletal reorganization, cell migration, and cancer metastasis. BioMed Res. Int. 2015, 409245 (2015).

62. Song, Y. et al. Mechanosensitive channel Piezo1 induces cell apoptosis in pancreatic cancer by ultrasound with microbubbles. iScience 25, 103733 (2022).

63. Gudipaty, S. A. et al. Mechanical stretch triggers rapid epithelial cell division through Piezo1. Nature 543, 118–121 (2017).

64. Kumar, N. et al. Piezo regulates epithelial topology and promotes precision in organ size control. 2023.08.16.553584 Preprint at 10.1101/2023.08.16.553584 (2023).

65. Pardo-Pastor, C. et al. Piezo2 channel regulates RhoA and actin cytoskeleton to promote cell mechanobiological responses. Proc. Natl. Acad. Sci. 115, 1925–1930 (2018).

66. Schvartzman, M. et al. Nanolithographic Control of the Spatial Organization of Cellular Adhesion Receptors at the Single-Molecule Level. Nano Lett. 11, 1306–1312 (2011).

67. Coyer, S. R. et al. Nanopatterning reveals an ECM area threshold for focal adhesion assembly and force transmission that is regulated by integrin activation and cytoskeleton tension. J. Cell Sci. 125, 5110–5123 (2012).

68. Roca-Cusachs, P., Iskratsch, T. & Sheetz, M. P. Finding the weakest link: exploring integrin-mediated mechanical molecular pathways. J. Cell Sci. 125, 3025–3038 (2012).

69. Takuwa, Y. et al. Calcium-dependent regulation of Rho and myosin phosphatase in vascular smooth muscle. Biomed. Rev. 16, 13–21 (2005).

70. Velagala, V. & Zartman, J. J. Pinching and pushing: fold formation in the Drosophila dorsal epidermis. Biophys. J. 120, 4202–4213 (2021).

71. Velagala, V., Chen, W., Alber, M. & Zartman, J. J. Chapter 4.1 - Multiscale Models Coupling Chemical Signaling and Mechanical Properties for Studying Tissue Growth. in *Mechanobiology* (ed. Niebur>, G. L.) 173–195 (Elsevier, 2020). doi:10.1016/B978-0-12-817931-4.00010-8.

72. Banwarth-Kuhn, M. et al. Combined computational modeling and experimental analysis integrating chemical and mechanical signals suggests possible mechanism of shoot meristem maintenance. PLOS Comput. Biol. 18, e1010199 (2022).

73. Nematbakhsh, A. et al. Multi-scale computational study of the mechanical regulation of cell mitotic rounding in epithelia. PLOS Comput. Biol. 13, e1005533 (2017).

74. Ramezani, A. et al. A multiscale chemical-mechanical model predicts impact of morphogen spreading on tissue growth. Npj Syst. Biol. Appl. 9, 1–12 (2023).

75. Brodskiy, P. A., et al. QuickStitch for seamless stitching of confocal mosaics through high-pass filtering and recursive normalization. http://biorxiv.org/lookup/doi/10.1101/075440<x> (2016) doi:10.1101/075440.

