## Supplementary Information for "Reverse engineering morphogenesis through Bayesian optimization of physics-based models"

### Supplementary Tables

**Table S1:** Table corresponding to Figure 4C

| $log(k_A^{col})$ | $log(k_B^{col})$ | $log(k_L^{col})$ | $log(L0_A^{col})$ | $log(L0_B^{col})$ | $log(L0_L^{col})$ | $log(K_{ECM})$ | $F_B$ | Group |
|---|---|---|---|---|---|---|---|---|
| -0.1457 | -6.4092 | 1.7781 | -2.7738 | -1.5063 | -0.0522 | -11.2278 | 0.09940 | Control-1 |
| -0.1457 | -6.4092 | 1.7781 | -2.7738 | -1.5063 | -0.0522 | -11.2278 | 0.0994 | Control-1 |
| -0.551 | -3.1890 | -1.7789 | -2.9772 | -1.3214 | -0.0258 | -13.1908 | 0.1042 | Control-1 |
| 0.1594 | -5.3218 | -0.0223 | -2.8371 | -1.4229 | -0.1745 | -15.2949 | 0.1083 | Control-1 |
| 0.0510 | -4.1404 | -3.3223 | -2.9066 | -1.8135 | -0.3121 | -15.2864 | 0.1096 | Control-1 |
| 0.0510 | -4.1404 | -3.3223 | -2.9066 | -1.8135 | -0.3121 | -15.2864 | 0.1096 | Control-1 |
| -0.3332 | -5.2156 | -0.9684 | -2.9125 | -1.4194 | 0.00271 | -7.5177 | 0.1153 | Control-1 |
| -0.2216 | -4.7614 | -5.2241 | -2.8923 | -1.2269 | -0.5623 | -16.6767 | 0.1165 | Control-1 |
| 0.2265 | -5.4243 | -0.3725 | -2.8076 | -2.2940 | -0.0523 | -14.9203 | 0.1166 | Control-1 |
| -7.4342 | 1.5032 | -1.6709 | -1.2590 | -1.7627 | 0.7887 | -13.9026 | 0.1113 | Control-2 |
| -7.4342 | 1.5032 | -1.6709 | -1.2590 | -1.7627 | 0.7887 | -13.9026 | 0.1113 | Control-2 |
| -8.27186 | 2.1807 | -0.5177 | -1.3336 | -1.7698 | 0.6753 | -18.1196 | 0.1122 | Control-2 |
| -11.2967 | -8.6174 | -3.9609 | -2.2909 | -2.4713 | 0.8820 | -17.9534 | 0.2348 | Clgns |
| -11.2967 | -8.6174 | -3.9609 | -2.2909 | -2.4713 | 0.8820 | -17.9534 | 0.2348 | Clgns |
| -8.87599 | -6.75723 | -5.89843 | -2.6936 | -1.6007 | 0.6700 | -7.1689 | 0.2386 | Clgns |
| -11.2266 | -6.86542 | -7.56488 | -2.2389 | -2.1785 | 0.9723 | -11.9379 | 0.2390 | Clgns |
| -11.2266 | -6.86542 | -7.56488 | -2.2389 | -2.1785 | 0.9723 | -11.9379 | 0.2390 | Clgns |

**Table S2:** Table corresponding to Figure 5C

| $log(k_A^{col})$ | $log(k_B^{col})$ | $log(k_L^{col})$ | $log(L0_A^{col})$ | $log(L0_B^{col})$ | $log(L0_L^{col})$ | $log(K_{ECM})$ | $F_B$ | **Group** |
|---|---|---|---|---|---|---|---|---|
| -8.3067 | -3.3898 | -7.1575 | -1.707 | -1.6217 | 0.8230 | -8.0974 | 0.1045 | Case I |
| -8.3097 | -11.2236 | -11.1142 | -1.7834 | -2.3492 | 0.0376 | -12.4807 | 0.1099 | Case I |
| -9.2125 | -0.4135 | 0.3028 | -1.8481 | -1.6338 | 0.7404 | -4.9901 | 0.1019 | Case I |
| -11.3945 | -0.4277 | -10.324 | -1.3742 | -1.3303 | 1.2532 | -4.7872 | 0.0824 | Case II |
| -10.3464 | 2.0910 | -4.4691 | -1.5400 | -1.6170 | 1.1392 | -5.1180 | 0.0818 | Case II |
| -8.8677 | 2.1366 | -2.0050 | -1.8368 | -1.4873 | 1.1661 | -6.0692 | 0.0891 | Case II |
| -10.5006 | 1.8216 | -2.2324 | -1.9311 | -1.4193 | 1.1403 | -5.9649 | 0.0821 | Case II |
| -9.7547 | -0.4106 | -0.4306 | -1.5697 | -1.7461 | 0.9965 | -5.192 | 0.0693 | Case II |
| -3.6974 | 1.5870 | -7.4071 | -1.8436 | -2.2116 | 1.1328 | -15.5845 | 0.1209 | Case III |
| -6.5249 | 1.1620 | -7.3866 | -1.5486 | -2.3996 | 1.2288 | -18.2644 | 0.1140 | Case III |
| -7.7476 | 0.1057 | -6.9335 | -2.8963 | -2.8410 | 1.2445 | -13.6869 | 0.1162 | Case III |
| -9.3780 | 1.5621 | -3.8466 | -1.2561 | -2.9289 | 0.9043 | -17.9181 | 0.1357 | Case IV |
| -8.9513 | 1.7643 | -0.9020 | -1.8360 | -2.9622 | 0.9749 | -13.3347 | 0.1197 | Case IV |
| -8.9194 | 1.9544 | -3.7264 | -1.5244 | -2.9255 | 0.9263 | -13.3634 | 0.1360 | Case IV |
| -9.9168 | 2.0507 | -1.7202 | -2.2472 | -2.8870 | 0.9852 | -18.2693 | 0.1197 | Case IV |

### Supplementary Figures

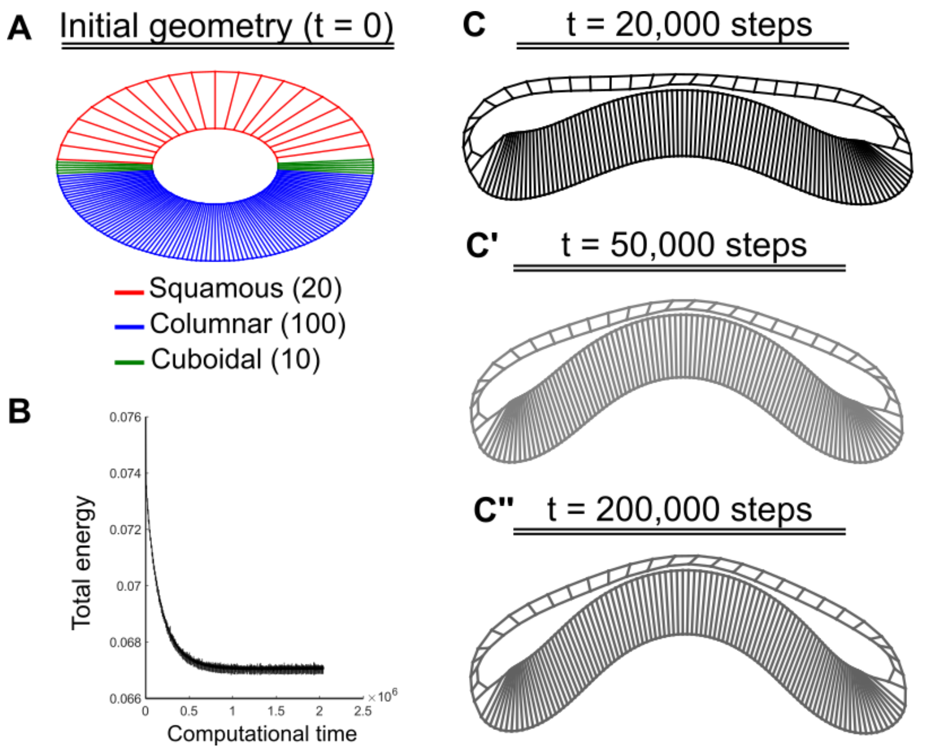


**Figure S1: Convergence of Surface Evolver simulations. (A)** The initial geometry used for modeling the AP cross-section of the wing disc resembles a donut. Parameters are specified differently for the different cell types named squamous, columnar, and cuboidal. **(B)** Plot showing variation in total energy with computational time. The energy reduces initially until it converges. **(C-C'')** Shape changes in the simulation cross-section with computational time. With energy, the shape also converges leading to a local minimum configuration.

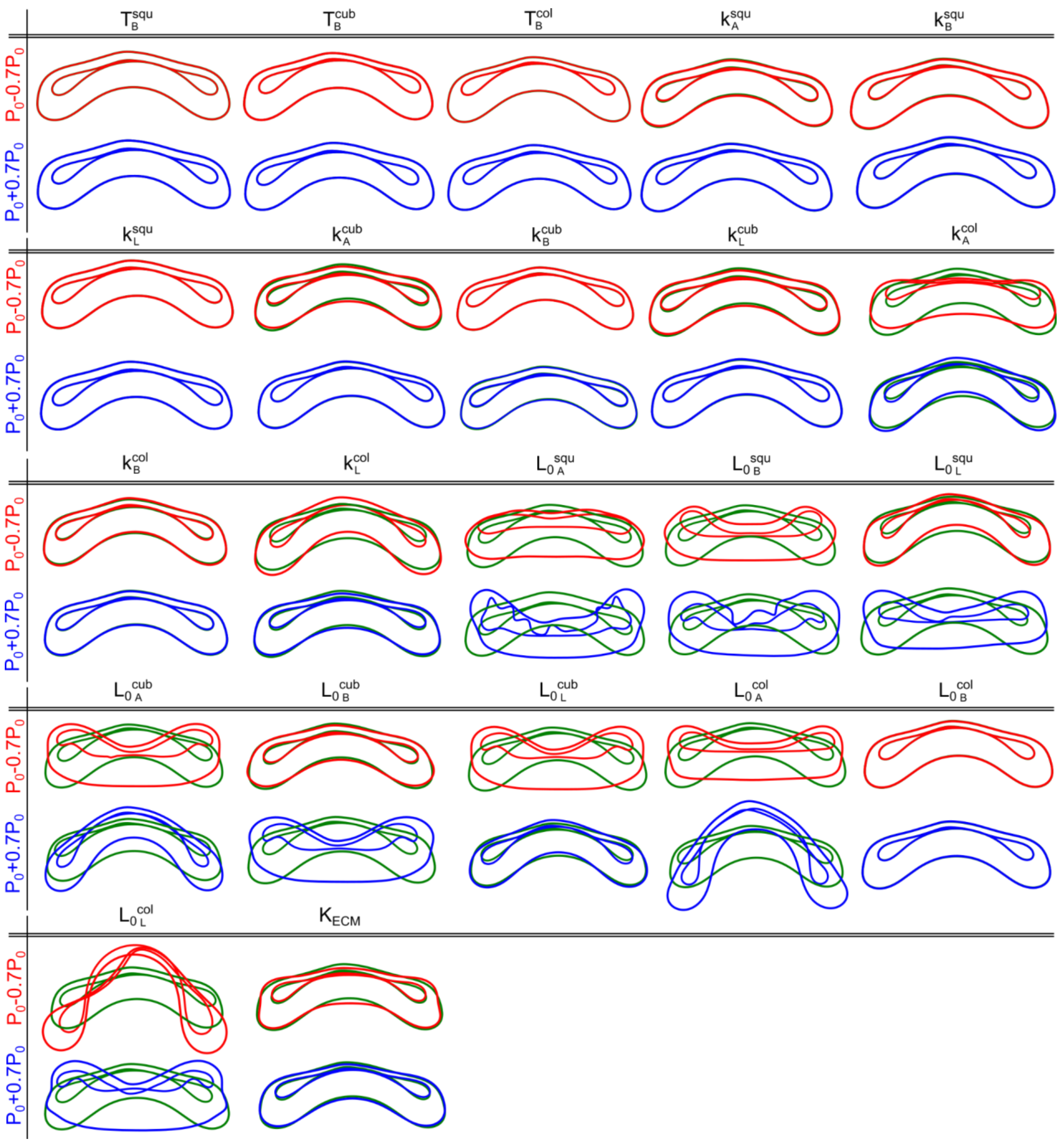


**Figure S2: Shape changes in wing imaginal disc as a result of perturbing individual parameters of the Surface Evolver model one at a time.** Individual model parameters within the model were increased and decreased by 70%, and Surface Evolver was run to measure the changes in tissue shape as a result of perturbation. The green contours within the plots indicate the base case. Blue and Red contours refer to the cases where the model parameters were increased and decreased respectively.

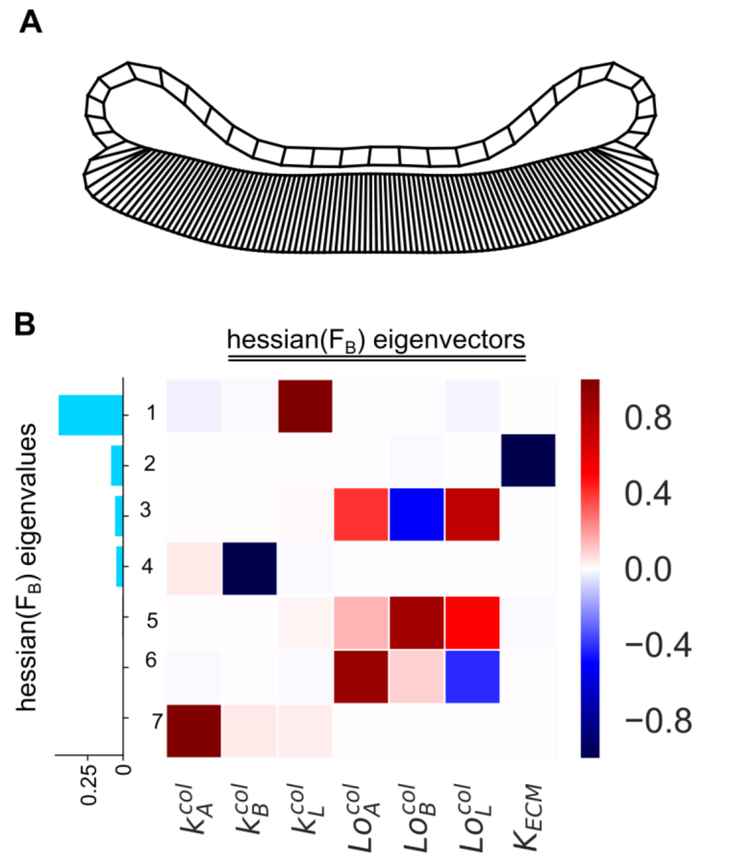


**Figure S3:** (A) Surface Evolver was used to simulate the minimum energy configuration for the average value of the parameters sampled (Table S1, Case: Clgns) using BO corresponding to the input wing disc cross-section shown in Figure 4A' (B) Eigendecomposition analysis of the local Hessian of $F_B$. Each row of the heatmap corresponds to the eigenvector of the Hessian matrix. The rows are arranged in decreasing order of the corresponding eigenvalues represented by a bar plot on the left-hand side of the heatmap.

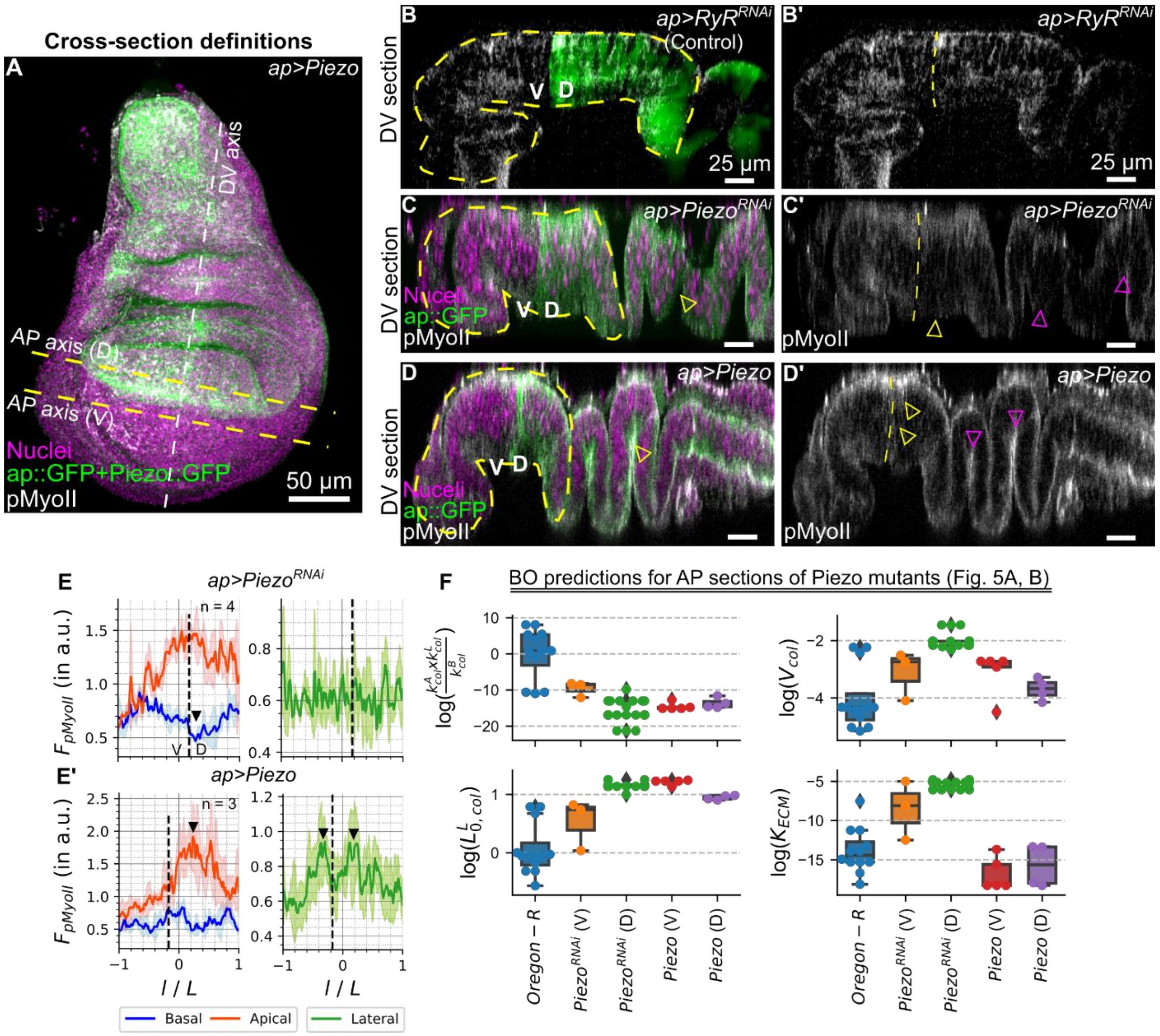


**Figure S4: Piezo regulates pMyoII expression during wing imaginal disc development.** An ap-Gal4 driver was used to downregulate and overexpress Piezo in the dorsal compartment of the wing disc using *UAS-Piezo$^{RNAi}$* and *UAS-Piezo:GFP* (BDRC# 58772) transgenic fly lines respectively. **(A)** A maximum intensity z-plane projection of the disc expressing *ap>Piezo*. Different lines within the pouch indicate the approximate location of the cross-section analyzed in Figure 5. **(B-B')** DV cross-section of the disc expressing ap>RyR$^{RNAi}$ (BDRC# 31540). ap>RyR$^{RNAi}$ has been used as a control for generating mutants as Ryanodine receptor (RyR) is not known to be expressed in the wing imaginal disc tissue[1] **(C-C')** DV cross-section of the disc expressing ap>Piezo$^{RNAi}$ (VDRC# 105132). **(D-D')** DV cross-section of the disc expressing ap>Piezo:GFP (BDRC# 58772). Fluorescent labels have been indicated within the plot. **(E,E')** Quantifying normalized fluorescent intensity of pMyoII along the apical-basal and lateral surfaces of the pouch. The x-axis of the pouch represents the normalized distance from the pouch center where the fluorescent intensity was calculated. Normalization of fluorescent intensity has been carried out by dividing the apical-basal signals with the max reported intensity value in the basal surface. A solid colored line represents the mean, and the shaded region represents the standard deviation in prediction from multiple samples. **(F)** Box plot showing the variation of parameter combinations (x-axis) for the best shapes predicted by the BO framework for cross sections presented in Figure 5. Parameters of the best shape predictions for the AP section of Oregon-R disc shown in Figure 4A have been used as a control for comparison.

### References

1. Consortium, T. modENCODE *et al.* Identification of Functional Elements and Regulatory Circuits by Drosophila modENCODE. *Science* **330**, 1787–1797 (2010).
